# A calibration framework to improve mechanistic forecasts with hybrid dynamic models

**DOI:** 10.1101/2022.07.25.501365

**Authors:** Victor Boussange, Pau Vilimelis Aceituno, Frank Schäfer, Loïc Pellissier

## Abstract

Process-based, dynamic models are essential for extrapolating beyond current trends and anticipating biodiversity responses to global change. However, their practical adoption for forecasting purposes remains limited due to difficulties in calibrating them against data and structural inaccuracies in their mathematical formulations. While ecological time series could, in principle, be used to directly estimate model parameters and refine model structures, the large noise levels in ecological datasets and the strong nonlinearity of ecological dynamics challenge conventional calibration methods. Here, we present a robust and scalable calibration framework that addresses these challenges by integrating techniques from scientific computing and deep learning. Our approach combines a segmentation strategy where state variables are estimated independently, differentiable programming for efficient gradient computation, parameter transformations to ensure the feasibility and stability of the model simulations, and mini-batching to accomodate large datasets. Through comprehensive benchmarks using simulated food web dynamics of increasing complexity, we demonstrate that the framework substantially improves the convergence of gradient descent algorithms and Monte Carlo sampling methods, accommodating for realistic scenarios with noisy and partial observations. This yields improved parameter estimation and forecasts within both mode estimation and full posterior distribution contexts. Crucially, we show that the calibration framework scales effectively with both the number of parameters and state variables. The improved convergence and scalability of the calibration framework enables hybrid modeling approaches, where neural networks parameterize complex processes within the dynamic model. In particular, we demonstrate that neural networks can effectively capture environmental dependencies in demographic rates and recover functional responses governing trophic interactions. Neural network-based parameterizations have the capability to improve the structural inaccuracies of models while maintaining ecological interpretability through post-hoc analysis of learned representations. We provide an implementation of the calibration framework and other key utilities as the open-source Julia package HybridDynamicModels.jl, with the hope that the package will facilitate the development of hybrid modeling approaches in quantitative ecology and related fields.

## 1 Introduction

Ecosystems are undergoing intense disruptions due to global changes [IPBES, 2019], which require models capable of extrapolating ecological dynamics beyond observed trends [Boyd, 2012]. A major challenge is that the processes governing ecological dynamics are nonlinear, leading to complex responses and feedback mechanisms [Scheffer et al., 2001]. Nonlinearity significantly limits the ability of modeling approaches that do not incorporate explicit biological knowledge to reliably project current trends into the future [Barnosky et al., 2012]. For example, data-driven methods such as species distribution models [Guisan and Zimmermann, 2000, Deneu et al., 2021] and nonparametric approaches for ecological time series prediction [Ye et al., 2015, Ye and Sugihara, 2016, Deyle et al., 2016] exhibit strong interpolation capabilities but are poorly suited for extrapolating into new collinearity structures [Barnosky et al., 2012, Urban et al., 2016]. In contrast, mechanistic ecosystem models impose constraints on expected dynamics by explicitly modeling interactions, feedback loops, and dependencies between ecosystem components [Geary et al., 2020]. Although such models are theoretically more robust for forecasting under large disturbances [Norberg et al., 2012], modelers often struggle fitting them against observational data [Fordham et al., 2018]. Difficulties arise primarily from two sources: inaccurate prior parameter values, which strongly influence calibration outcomes, and inaccuracies in the mathematical formulation of ecological processes [Schartau et al., 2017, Cabral et al., 2017, Briscoe et al., 2019, Pagel and Schurr, 2012]. To overcome these limitations and improve predictions of ecosystem responses to global changes, novel approaches that combine the inductive bias of mechanistic models with observational data are needed [Hartig et al., 2012].

Dynamic models can be calibrated against data by defining an inverse problem, where model parameter estimates are refined through inference methods by matching the model outputs to the observations. Inference methods can be broadly classified as either gradient-based or gradient-free approaches and as mode estimation or full posterior distribution estimation methods (see Box 1 for details). The computational complexity of gradient-based methods scales better with the number of parameters and state variables than gradient-free approaches. mode estimation methods naturally require significantly fewer model simulations compared to full posterior distribution estimation methods, but the latter allow for characterizing parameter and forecast uncertainties. Both mode and full posterior estimation methods face critical challenges in the context of dynamic ecosystem models. First, these methods require repeated model simulations, yet dynamic models are computationally expensive, limiting the number of feasible iterations [Fisher et al., 2018]. Second, the complexity of ecological processes often requires a large number of parameters to characterize their influence on state variables, while the curse of dimensionality [Boyd, 2012] makes posterior exploration increasingly more difficult as the number of parameters increases [Gutenkunst et al., 2007]. Third, available time series are often sparse, noisy, and incomplete with partially observable state variables [Scheiter et al., 2013, Schartau et al., 2017], diverging from traditional inverse problem settings. Ecosystem dynamics are also highly sensitive to initial conditions (ICs) and may exhibit chaotic behavior [Hastings et al., 1993, Huisman and Weissing, 1999, Benincà et al., 2008], or show a wide range of possible dynamics depending on parameter values. In such cases, small perturbations in IC or parameter values can lead to large divergences in model outcomes, leading to convergence issues during the calibration process [DeAngelis and Yurek, 2015, Pagel and Schurr, 2012, Vilimelis Aceituno et al., 2025].

Beyond the challenges specific to inference methods, the mathematical representation of ecosystem processes is inherently difficult due to their complexity, resulting in ecosystem models that show large structural inaccuracies [Wood, 2001, Gehlen et al., 2015, Schartau et al., 2017, Purves et al., 2013]. For instance, functional responses, which describe how resource intake varies with resource density [Rosenbaum and Rall, 2018], as well as the dependence of demographic parameters on species traits [Chalmandrier et al., 2022] or environmental drivers [Amarasekare and Johnson, 2017], are often inadequately represented in mechanistic models. These inaccuracies can lead to erroneous predictions and spurious dynamics [Gentleman et al., 2003]. This challenge extends beyond ecology to other natural sciences, where neural networks have been used successfully to parameterize complex processes in weather and climate systems [Kashinath et al., 2021, Peng et al., 2021, Rasp et al., 2018], glacier dynamics [Bolibar et al., 2023], and systems biology [Alber et al., 2019], leading to substantial improvements in predictive performance [Willard et al., 2020]. Neural network-based parameterization of ecological processes holds similar promise for ecosystem models, but they represent additional challenges for inference methods, as they contribute to substantially increasing the dimensionality of the parameter space. The integration of neural network-based parameterization in dynamic ecosystem models requires robust and scalable calibration methods that accommodate the noisy nature of ecological time series.

Multiple shooting methods, also known as segmentation strategies or piecewise regression, have been developed to improve the robustness of calibration methods when applied in the context of dynamic models Pisarenko and Sornette [2004], Hamilton [2011]. These techniques partition the time series into short segments; by matching model simulations against these short segments, the high sensitivity of the model to parameters is mitigated, which in turn smooths the posterior distribution and improves the convergence of the calibration method [Vilimelis Aceituno et al., 2025]. We provide an illustration of this effect in Fig. 1. However, segmentation strategies are mostly used together with fixed initial conditions [Hamilton, 2011], which does not suit the noisy and partially observable nature of ecological time series. Moreover, the proposed methodologies do not take advantage of recent advances in differentiable programming [Sapienza et al., 2024] and scalable optimization routines that have been developed in the context of deep learning.

**Figure 1.**
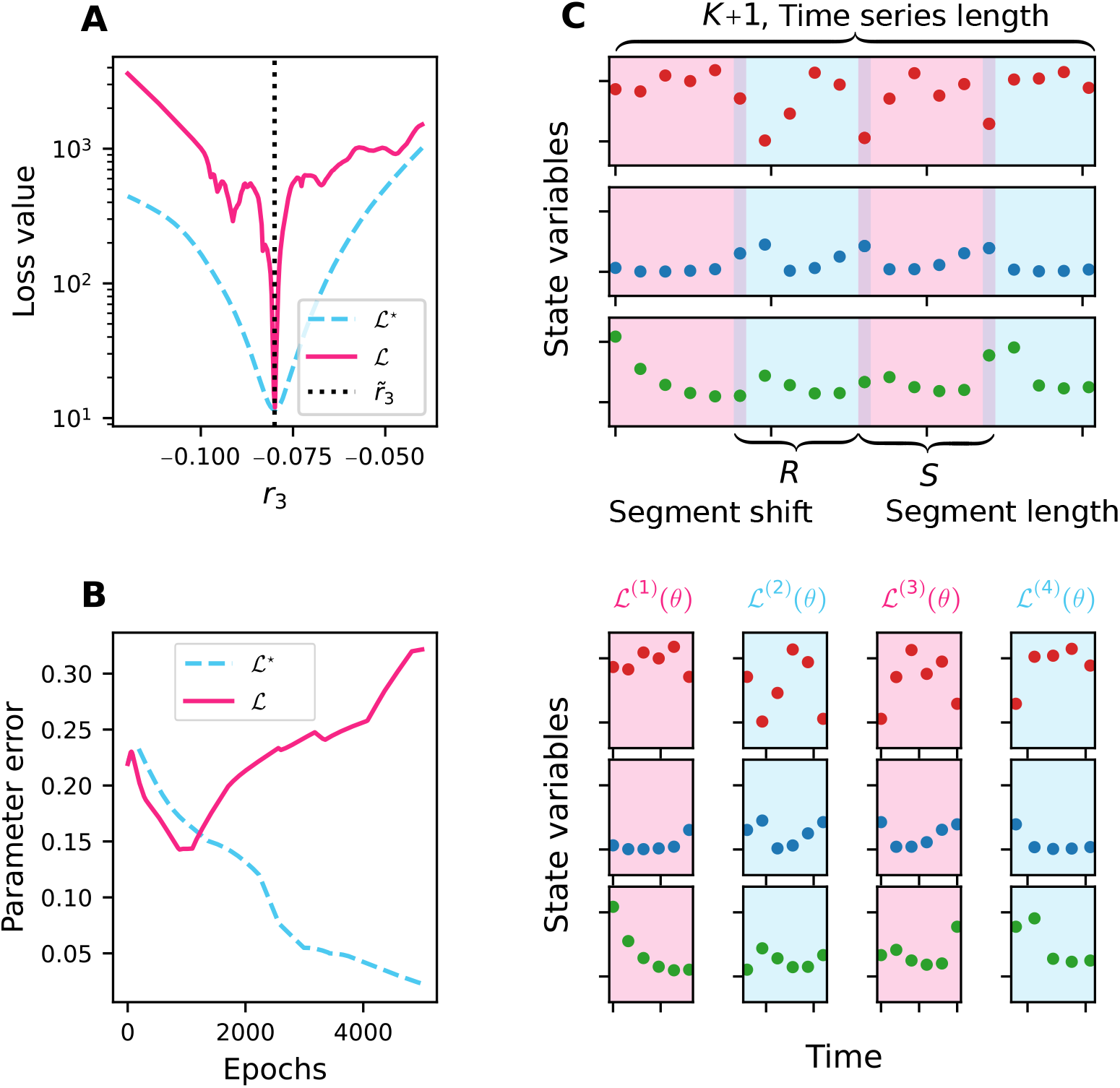
Illustration of the segmentation method. **A**. Cross-section of the loss surfaces defined by the naive loss function ℒ (red plain curve) and the piecewise loss function ℒ ^⋆^ (blue dashed curve), for model ℳ_3_ (see Section 3 for details). The surface of ℒ exhibits multiple local minima and steep gradients near the true parameters, making gradient descent-based optimization difficult. In contrast, ℒ^⋆^ is smooth and has a single minimum, demonstrating the regularization effect of partitioning the time series into shorter segments. **B**. Convergence of the Adam optimizer when minimizing ℒ (red plain curve) and ℒ^⋆^ (blue dashed curve) for model ℳ_3_. The red curve diverges from the true parameter values after approximately 1000 iterations, while the blue dashed curve exhibits smooth convergence. **C**. Graphical representation of the proposed segmentation method. The algorithm partitions the time series into *M* segments of *S* data points (blue and red segments) with shift *R*.

Here, we present a comprehensive calibration framework designed to address the unique challenges posed by mechanistic ecological forecasting. Our framework integrates the partitioning of the available time series into segments, where initial conditions are estimated independently. We constrain model parameter values within feasible ranges through parameter transformation, and leverage differentiable programming to efficiently obtain the gradient of the model with respect to its parameters. By conducting a grid search over the hyperparameters of the method across simulated food web dynamics, we demonstrate the benefits of the proposed framework both within a mode estimation context with gradient descent optimization and a full posterior distribution estimation context with Monte Carlo sampling. We show that the calibration framework improves the estimation of parameters and forecasts over naive segmentation strategies, is robust against noisy and partial observations, and scales well with model complexity. These desirable properties make the framework instrumental for hybrid modeling. We present applications of hybrid modeling in two realistic settings, showing how neural networks can be used to parameterize complex ecological processes.

### Box 1. Overview of calibration methods

Calibration methods can typically be classified as gradient-free or gradient-based methods, and can be used to compute a point estimate of the parameters (e.g., the maximum likelihood estimate or maximum a posteriori estimate, see Box 2) or estimate the full posterior distribution. Bayesian inference with Monte Carlo sampling, as applied in Lignell et al. [2013], Higgins et al. [2010], Xu et al. [2006], Fiechter et al. [2013], Rosenbaum et al. [2019], Heiland et al. [2023], Rosenbaum and Fronhofer [2023], Pagel and Schurr [2012], enables the quantification of uncertainties by estimating the full posterior probability distribution of the parameters. This is achieved through a global exploration of the parameter space, but it comes at a high computational cost. Monte Carlo sampling methods are particularly vulnerable to the curse of dimensionality [Gosh et al., 2021], where the volume of the parameter space grows exponentially with the number of parameters, necessitating increased sampling. Hamiltonian Monte Carlo sampling methods improve sampling efficiency by leveraging gradient information [Hoffman and Gelman, 2011], yet they remain computationally more expensive than point estimation approaches. Simulated annealing [Matear, 1995] and genetic algorithms [Ward et al., 2010] are gradient-free point estimation methods that do not rely on posterior gradients. As with Monte Carlo approaches, they suffer from the curse of dimensionality and require a large number of model evaluations. Gradient-based point estimation methods, such as gradient descent algorithms (see Box 2), are typically the most computationally efficient, requiring significantly fewer model evaluations [Schartau et al., 2017]. This efficiency explains their widespread adoption in the calibration of marine ecosystem models [Fennel et al., 2001, Spitz et al., 1998, Xiao and Friedrichs, 2014, Pelc et al., 2012] (see Schartau et al. [2017] for a review) or terrestrial ecosystem models [Zhu and Zhuang, 2015, DeLong et al., 2014, Curtsdotter et al., 2019], and their wide adoption in deep learning for training highly parameterized models.

## 2 Materials and methods

### 2.1 Ill-conditioning of the inverse problem

The calibration of dynamic models in a point estimation context can be formulated as an optimization problem where the goal is to minimize a loss function ℒ associated with the posterior distribution (see Box 2 for details). While gradient descent algorithms generally offer computational advantages due to their favorable scaling properties, they frequently fail to converge when naively used to calibrate dynamic ecosystem models due to fundamental properties of ecological dynamics that create ill-conditioned inverse problems.

#### Box 2. Gradient descent for the calibration of dynamic models

**Dynamic models**

Deterministic ecosystem models typically consist of a system of ordinary differential equations (ODEs) describing the evolution of state variables (e.g., species abundances, resource availability) over time:

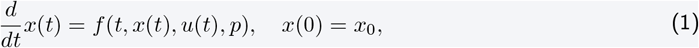

where *x*(*t*) ∈ ℝ^*m*^ represents the state variables at time *t, u*(*t*) denotes external forcings, *p* ∈ ℝ^*q*^ is the parameter vector, and *x*_0_ ∈ ℝ^*m*^ are the initial conditions. The model solution can be expressed as a mapping ℳ (*t, θ*) = *x*(*t*) where *θ* = (*x*_0_, *p*) ∈ ℝ^*m*+*q*^ is a parameter vector combining initial conditions and parameters.

**Observation model**

It is general practice to assume that observations 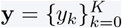 at discrete time points *t*_0_, *t*_1_, …, *t*_*K*_ are related to the true state variables through an observation model. Let *h* : ℝ^*m*^ →ℝ^*d*^ be an observation function that maps the full state space to the observation space. The observation model consists of the probability distribution whose probability density function *f*_*x,σ*_ is parameterized by the state variables and the observation error parameters *σ*. This formulation accommodates partial observations (when *d < m*) and various noise structures.

**Maximum a posteriori estimation with gradient descent**

Given observations **y** and assuming independence across time points, the likelihood function may be expressed as:

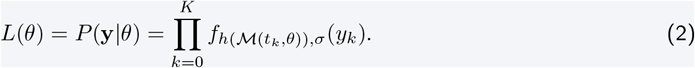

Incorporating prior knowledge *P* (*θ*) about parameters and initial conditions, Bayes’ theorem gives the posterior distribution:

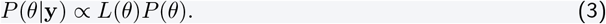

The maximum a posteriori (MAP) estimator maximizes the posterior distribution:

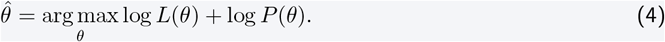

When the prior *P* (*θ*) is uniform (uninformative), MAP estimation reduces to maximum likelihood estimation (MLE), where 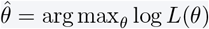. Equation (4) can be expressed as an equivalent optimization problem where the objective is to minimize a loss function ℒ which generally takes the form:

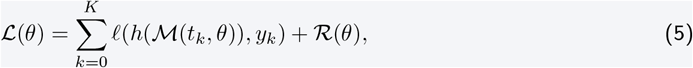

where *ℓ* defines a discrepancy measure between the model predictions and the observations, and ℛ (*θ*) is a regularization term. For example, assuming log-normally distributed observations *y*_*k*_ ∼ LogNormal(*x*(*t*_*k*_), Σ_*y*_) and Gaussian priors *θ ∼* 𝒩 (*θ*_*b*_, Σ_*p*_), the optimization problem in Eq. (4) boils down to minimizing the loss function:

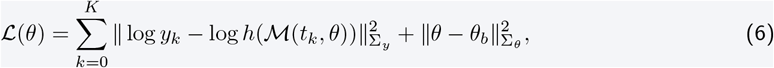

where 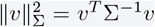 denotes the Mahalanobis distance. The second term can be viewed as a L2 regularization term, with the regularization strength determined by the prior covariance Σ_*θ*_.

Gradient descent permits to minimize ℒ by iteratively updating the parameter vector *θ* using the gradient of the loss function:

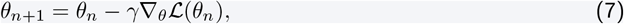

where *γ >* 0 is the learning rate and *∇* _*θ*_ℒ is the gradient with respect to *θ*. Modern gradient descent algorithms incorporate more advanced update strategies, such as adaptive learning rates and momentum, for improved convergence.

The loss function associated with nonlinear ecological dynamics generally exhibits numerous local minima that prevent gradient descent algorithms from reaching the global minimum (Fig. 1**A**-**B**, red curve). In addition, it presents steep gradients near the true parameters, causing numerical overflows and leading gradient descent algorithms to overshoot optimal solutions. These properties arise from the sensitivity of dynamic ecosystem models to small perturbations in parameters and initial conditions (ICs). [Vilimelis Aceituno et al., 2025] formally shows that the higher this sensitivity, the larger the number of local minima in the loss function, and the steeper the gradients near true parameter values. The sensitivity of dynamic models is particularly pronounced in ecosystem models, which often show chaotic or cyclic behavior [Bjørnstad and Grenfell, 2001, Hastings et al., 1993, Huisman and Weissing, 1999, Benincà et al., 2008]. Our empirical results in Section 3.2 confirm that gradient descent algorithms consistently converge to suboptimal local minima when applied to the standard loss formulation (Eq. (6)). Although techniques such as adaptive learning rates, gradient normalization, and learning rate schedulers can partially mitigate issues associated with excessively large gradients near the true parameters, they cannot address the fundamental problem of multiple local minima that severely limits the convergence of gradient descent algorithms. The ruggedness of the loss function directly translates to the posterior distribution, creating analogous difficulties in a full posterior estimation context with Monte Carlo sampling methods, as we show in Section 3.2.

Beyond the ill-conditioning of the posterior distribution, one has to carefully safeguard against parameter combinations resulting in biologically infeasible model simulations, with e.g. solutions to the ODE system diverging to infinity, which may destabilize the entire calibration process. Out-of-the box gradient descent optimization algorithms do not allow for a suitable formulation of the feasible region.

We address both the convergence and the numerical instability issues through an improved formulation of the inverse problem, which we present in the following section.

### 2.2 A robust and scalable calibration framework for dynamic models

#### 2.2.1 Description of the calibration framework

We present a comprehensive calibration framework that addresses the convergence and numerical stability issues identified in the previous section. Our approach integrates four key components: (i) a segmentation strategy with independent initial condition estimation, (ii) parameter transformations that constrain values to both numerically and biologically feasible ranges, (iii) mini-batching to reduce memory requirements, and (iv) differentiable programming for efficient gradient computation.

##### Time series segmentation and initial condition estimation

Given a time series with *K* + 1 observations **y**_0:*K*_, we partition it into segments of length *S*, shifted by *R* data points (see Fig. 1**C** for an illustration). Assuming for simplicity that *K* + 1 is a multiple of *S* − *R*, this results in 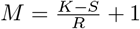 segments:

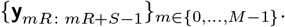

The segment shift parameter *R* allows us to control the overlap between segments, with *R* ≥ *S* corresponding to non-overlapping segments and *R* = 1 to maximally overlapping segments. A large overlap increases the number of segments, which may improve parameter estimation accuracy but also increases computational cost. Setting *R > S* allows us to isolate held-out validation data points from the training data, which may be used to prevent overfitting; we use this strategy in Section 3.3.

Rather than fixing initial conditions at segment boundaries to observed values, we treat them as additional parameters to be estimated. This approach is essential for two key scenarios: first, when handling noisy data where observed values may deviate substantially from the true underlying state variables, and second, when accommodating partially observable systems where some state variables are unobserved and cannot be directly inferred from the available data.

##### Parameter transformations for numerical stability

To constrain parameter and state variable values to a feasible region of the parameter space, we use parameter transformations by defining the bijective maps 𝔓: 𝒫 ∋ *p* → 𝔭 ∈ ℝ^*q*^ and 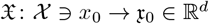 that allow for optimization in the unconstrained real domain ℝ^*q*^ × ℝ^*d*^ while guaranteeing that the inverse images 𝔓^−1^(𝔭) and 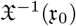 remain in the feasible domains *P* ⊂ ℝ^*q*^ and 𝒳 ⊂ ℝ^*d*^.

##### Piecewise loss function with mini-batching

We reformulate the inverse problem in Box 2 as a piecewise inverse problem that operates on mini-batches of segments. Given a random mini-batch 𝒮⊂ {0, …, *M* − 1} that selects |𝒮 | = *b* segments, the segmented loss function becomes:

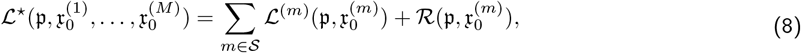

where

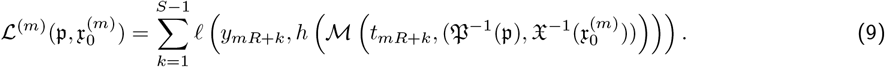

Here, 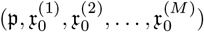 represents the augmented parameter vector in the unconstrained domain, containing *M* additional initial conditions 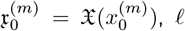 is a discrepancy measure between model predictions and observations (e.g., mean squared log error, see Box 2), and ℛ is a regularization term.

This reformulation of the loss function smooths its surface, improving the convergence of optimization or sampling methods (see Fig. 1**A**, blue dashed curve, for an illustration). In addition, this piecewise formulation also allows to correct for residual process error, similarly to state-space models [Auger-Méthé et al., 2021].

##### Gradient descent

Each optimization step consists of: (i) sampling a new mini-batch 𝒮, (ii) updating parameters 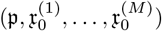 using a gradient descent algorithm, and (iii) repeating this process until all segments have been used (one epoch), then iterating over multiple epochs.

##### Bayesian inference with Monte Carlo sampling

The piecewise inverse problem can be analogously formulated in a full posterior distribution estimation context, where parameters and initial conditions for each segment are associated with a prior distribution (see Section 3 and numerical implementation in Section 8).

This framework introduces three key hyperparameters: the mini-batch size *b*, the segment size *S*, and the segment shift *R*. These parameters influence the parameter estimation accuracy, convergence, computational cost, and memory footprint. We provide a comprehensive analysis of their effects in Section 3, and demonstrate that an appropriate choice of these hyperparameters leads to substantially improved convergence of both gradient descent algorithms and Monte Carlo sampling. Despite the increased dimensionality of the parameter space resulting from inferring additional initial conditions, we also show that the combination of mini-batching and the use of appropriate sensitivity methods for ODEs ensures computational efficiency.

#### 2.2.2 Numerical implementation of the calibration framework

We develop a comprehensive software toolbox to facilitate the implementation of the calibration framework described above, released as the open-source Julia package HybridDynamicModels.jl. Built on top of the deep learning library Lux.jl[Pal, 2023], the package provides composable utility layers to easily specify dynamic models and implement the segmentation strategy. The core components of the package include: (i) ODEModellayers for defining differential equation systems, optionally constructed from combinations of other Lux.jllayers, (ii) ParameterLayers to specify learnable parameters with optional transformations via Constraintstructures, (iii) ICLayers to facilitate initial condition estimation for each segment, (iv) BayesianLayers to wrap any Lux.jllayer with parameter priors for full posterior distribution estimation, (v) utilities for time series segmentation and mini-batching, and (vi) a unified training API. The ODEModellayer relies on OrdinaryDiffEq.jl[Rackauckas and Nie, 2017] for efficient numerical integration, while gradients of the numerical solutions with respect to parameters and initial conditions are computed using custom differentiation rules from SciMLSensitivity.jl [Ma et al., 2021]. This package provides specialized algorithms that scale efficiently with both the number of state variables and parameters (see Sapienza et al. [2024] for details). Beyond the differential equation models which are the focus of this paper, HybridDynamicModels.jlalso supports autoregressive (ARModel) and closed-form models (AnalyticModel), that can be specified from combinations of Lux.jllayers. HybridDynamicModels.jlintegrates seamlessly with the broader Lux.jland Turing.jlecosystems. In particular, ODEModelcan be directly combined with neural network layers, enabling hybrid models where selected ecological processes are parameterized by neural networks (see Section 3.3). Our package naturally interfaces with the optimization libraries compatible with Lux.jl, allowing for the use of state-of-the-art gradient descent algorithms provided by the latter. The training API implements the segmentation strategy with initial condition inference as described in Eq. (8) through the trainfunction. For gradient-based optimization, the trainfunction works with the SGDBackendand is compatible with gradient descent algorithms from Optimisers.jl. To estimate the full posterior distributions, the trainfunction works with the MCSamplingBackend, enabling the use of various Monte Carlo sampling methods by interfacing with the Turing.jllibrary [Fjelde et al., 2025]. Complete documentation and tutorials are available at https://github.com/vboussange/HybridDynamicModels.jl. We use HybridDynamicModels.jlto implement all simulations presented in the following sections.

## 3 Experiments

### 3.1 Experimental set-up

We consider a generalized dynamic food-web model inspired by [Hastings and Powell, 1991, McCann and Yodzis, 1994b,a, Klebanoff and Hastings, 1994, Post et al., 2000, Åkesson et al., 2021], where species interact through trophic and competitive relationships. Details of the model formulation are provided in Section S1. We specialize the model into three variants with increasing complexity: ℳ_3_, ℳ_5_, and ℳ_7_, simulating the dynamics of 3, 5, and 7 species with 9, 18, and 24 parameters, respectively. The food-web structure of each ecosystem is illustrated in Fig. S1. These models have been used as benchmarks for ecosystem forecasting methods [Perretti et al., 2013, Deyle et al., 2016, Ye and Sugihara, 2016] and produce fluctuations that resemble observed ecological time series [Bjørnstad and Grenfell, 2001] while being challenging to forecast [Post et al., 2000]. We set the model parameters to biologically realistic values proposed by [McCann and Yodzis, 1994a, McCann and Hastings, 1997], ensuring that the dynamics of the system is chaotic or oscillatory (see Section S1 for numerical values of the parameters and Fig. S2 for an illustration of the dynamics). Time is scaled relative to the growth rate of the primary producer, so that its biomass increase is 100% per day. To generate observational data, we simulate ecosystem dynamics and introduce observation error by sampling from a log-normal distribution with mean equal to the true population size and covariance matrix Σ = *r*^2^*I*, where *r* represents the noise level. This observation model is commonly used in population dynamics [Schartau et al., 2017]. We exclude transient dynamics by sampling after a long burn-in period (*t >* 500, see Fig. S2). Following, we set *ℓ* to be the mean squared logarithmic error and employ uninformative parameter priors (the regularization term in Eq. (8) is set to 0). For each parameter *p*_*i*_, we randomly draw initial estimates from a uniform distribution with range 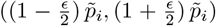, where 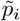 denotes the true generating parameter value and *ϵ* is a perturbation parameter. We constrain the parameter values to the same range and initial condition values to (10^−3^, 5). For gradient computation, we use the BacksolveAdjoint method from SciMLSensitivity.jl, which we found to be the most efficient. In the context of our calibration framework, this method has theoretical scaling 𝒪 (*b*(2*d* + *q*)) with *d* the number of state variables, *q* the number of parameters, and *b* the mini-batch size, thus scaling linearly with parameter dimensionality. For models with fewer parameters or state variables, forward sensitivity methods, with less favourable scaling ( 𝒪 (*bd*(*d* + *q*))) but less computational overhead, may be preferable [Ma et al., 2021, Sapienza et al., 2024]. We employ the Adam algorithm [Kingma and Ba, 2014] from Optimisers.jl with default parameters and a constant learning rate, setting the number of epochs to 3000. While Adam typically benefits from learning rate scheduling [Godbole et al., 2023], we use constant rates in the following section for the sake of simplicity.

### 3.2 Effect of segment length, initial condition estimation and batch size on parameter error, forecast error, and computational complexity

We demonstrate that a judicious choice of segment length combined with independent initial condition estimation consistently yields superior performance for both parameter accuracy and forecast skill across all models investigated, while adding reasonable computational overhead compared to naive gradient descent calibration strategies.

#### 3.2.1 Experimental design

We generate single time series for each reference model ( ℳ_3_, ℳ_5_, ℳ _7_), sampling every 4 days to obtain 111-point time series (Fig. 1**C**). The first 101 points serve as training data (*K* = 100), while the final 10 points are reserved as a test dataset for forecast evaluation.

We assess performance using two complementary metrics: (i) median relative parameter error across estimated parameters to evaluate parameter estimation, and (ii) forecast error calculated as the squared logarithmic difference between the last 10 observed points and predictions generated using estimated initial conditions from the final training segment. We systematically evaluate error statistics over 5 simulation runs.

#### 3.2.2 Optimal segment length and initial condition estimation

We conduct a grid search over varying noise level, segment lengths, and learning rates, with or without estimation of initial conditions, using mini-batch size *b* = min(10, *M* ) and shift *R* = *S* − 1 (overlap between segments of 1), and report results in Fig. 2. Using the full segment length (*S* = *K*, equivalent to the naive loss function ℒ, Eq. (6)), systematically results in poor performance regardless of learning rate. The smallest segment length (*S* = 2) also proves suboptimal, while intermediate lengths (*S* ∈ [4, 20]) yield best performance. This phenomenon arises because decreasing segment length flattens the loss surface, improving the convergence of gradient descent algorithms at the cost of a reduced curvature (Fig. 1**A**, Vilimelis Aceituno et al. [2025], Mikhaeil et al. [2022]). Consequently, excessive segmentation deteriorates parameter precision because the loss function exhibits similar values in extended neighborhoods around true parameters. The estimation of initial conditions at each segment proves crucial for achieving good performance, with all optimal hyperparameter combinations involving initial condition estimation. Estimating initial conditions becomes critical when data is highly noisy or in partial observation settings where some state variables are unobserved and initial conditions cannot be directly retrieved from data.

**Figure 2.**
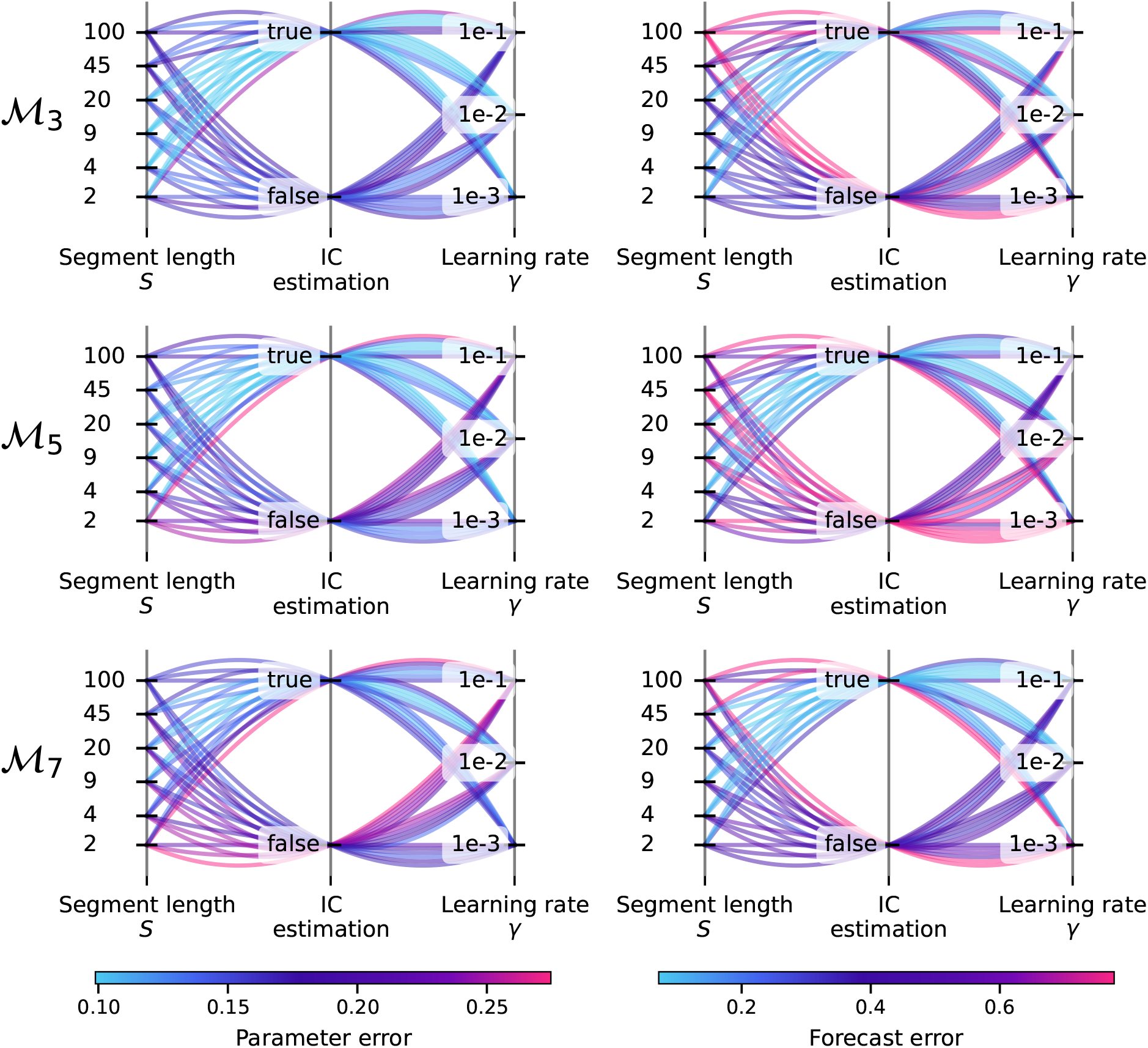
Parallel coordinates plot showing the effect of the combination of segment length *S*, estimation of ICs and learning rates on parameter error and forecast error, for varying model complexity. Panels along a row correspond to results for model ℳ_3_ (top, three state variables), ℳ_5_ (middle, five state variables) and ℳ_7_ (bottom, seven state variables), respectively. Panels along columns show the effects on parameter error (left) and forecast error (right), respectively. Results are obtained with a perturbation parameter *ϵ* = 1, a noise level *r* = 0.2 a fixed minibatch size *b* = min(10, *M* ) and a segment shift *R* = *S* − 1. Similar results obtained for higher noise level are reported in Fig. S3.

We find that these trends also hold for larger noise levels (Fig. S3) and in a full posterior distribution estimation setting using Hamiltonian Monte Carlo sampling (Fig. S4).

We emphasize that these results are specific to the system’s predictability horizon, to the model’s structural errors, and to the observation errors. As the level of noise in the observations decreases, the optimal segment length is likely to increase. In contrast, high stochasticity in the system may favor short segments to mitigate process errors (Turchin [2003], see Vilimelis Aceituno et al. [2025] for a discussion). In practice, the optimal hyperparameter combination for specific models and datasets can be determined by splitting the time series into a training and a validation dataset, and selecting hyperparameters based on the forecast skill obtained on the validation dataset (see Section 3.3, and Godbole et al. [2023] for in-depth recommendations on hyperparameter tuning).

#### 3.2.3 Scalability and computational considerations

The calibration framework scales effectively with model complexity, showing similar parameter and forecast error across all three models despite varying numbers of state variables and parameters (Fig. 2). This demonstrates that the benefits of piecewise formulation are maintained as ecological models increase in complexity. Our simulations show that the estimation of initial conditions introduces negligible computational overhead (blue vs red lines in Fig. 3), and that the computational cost of the framework scales well with model size (Fig. 3**C**). The segmentation strategy introduces computational overhead compared to the naive approach, as the number of ODE solves per epoch increases linearly with the number of segments (Fig. 3**A**). However, this overhead can be mitigated by increasing batch sizes (Fig. 3**B**) at the cost of increased memory requirements. In contrast, since Monte Carlo sampling methods scale poorly with parameter dimensionality, estimating full posterior distributions with these methods becomes computationally prohibitive for complex models or when inferring initial conditions at each segment (Fig. S4**C**). This limitation suggests that the inference of initial conditions should be mostly considered in the context of mode estimation with gradient descent algorithms, which should also be preferred for high-dimensional models. Monte Carlo sampling methods remain interesting options for simpler mechanistic models when uncertainty quantification is necessary.

**Figure 3.**
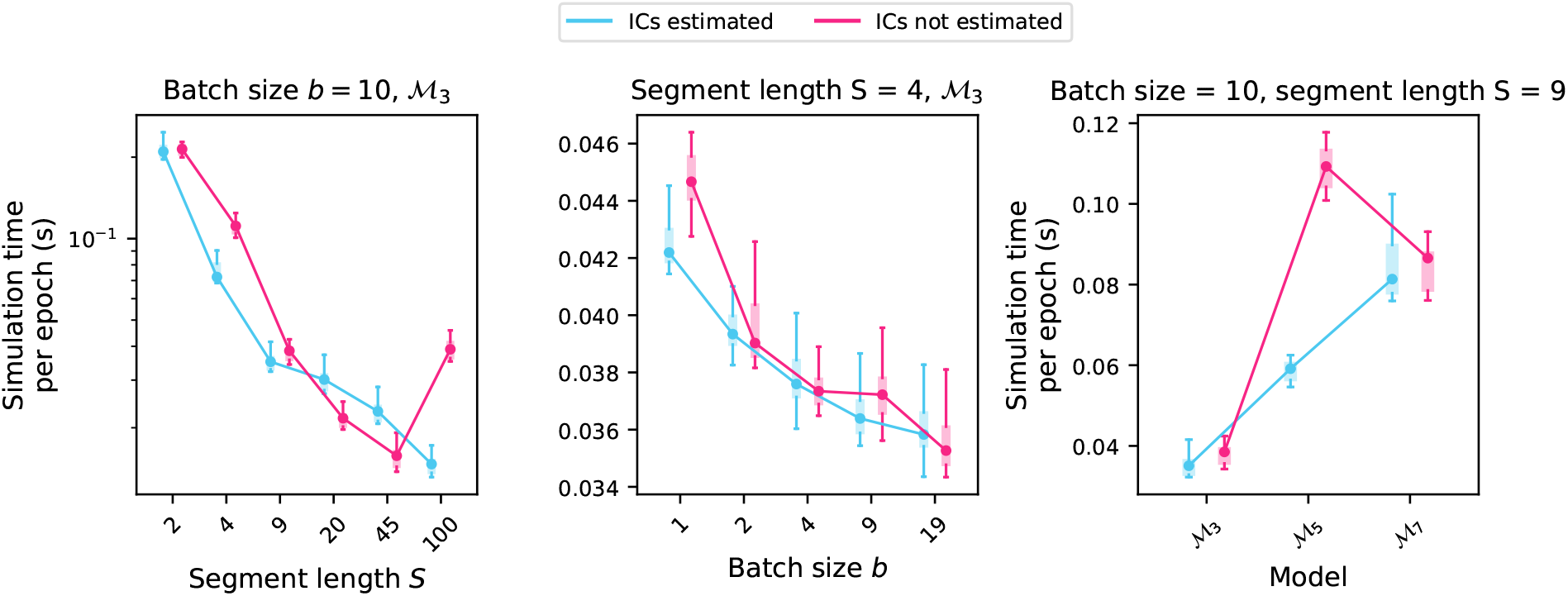
Effect of segment length *S*, batch size *b* and model complexity on simulation time for a single epoch. The panels show that reducing segment length increases simulation time, an effect that can be mitigated by increasing batch size. The overall framework scales well with model size, and estimating initial conditions introduces negligible computational overhead. Results are obtained with a segment shift *R* = *S* − 1.

### 3.3 Learning model parameterization with neural networks

The robustness and scalability of our calibration framework combined with gradient descent algorithms enables the parameterization of ecological processes using neural networks. We conduct two experiments demonstrating neural network parameterization of: (i) per-capita feeding rates of consumer and predator species, and (ii) growth rates of primary producer species. For both scenarios, we generate synthetic observations by adding noise (level *r* = 0.1) to reference model simulations. Following Section 3.2, we split datasets into training and test datasets and use inferred initial conditions from the final segment to generate forecasts. We evaluate forecast error statistics over 5 simulation runs. We set hyperparameters to *S* = 11, *R* = *S* + 1, and *b* = 10. By setting the segment shift greater than the segment length, we create held-out validation data points from the training set. This enables the computation of a validation loss to prevent overfitting, where we select the model parameters that minimize the validation loss across training epochs.

#### 3.3.1 Neural network-based parameterization of feeding rates

We attempt to recover functional responses of consumer and predator species using a hybrid model based on the structure of ℳ_3_, where nonzero coefficients of the per-capita feeding rate matrix are parameterized by a neural network. This network takes primary producer and consumer abundances as input and predicts feeding rates (Fig. 4**A**, details in Section S1.1.2). The neural network architecture consists of two input nodes, three hidden layers with *N*_*h*_ neurons each, and two output nodes, using hyperbolic tangent activation functions except in the final layer. We employ the Adam algorithm with decoupled weight decay regularization [Loshchilov and Hutter, 2019] (AdamWfrom Optimisers.jl), with weight decay parameter *λ* and other hyperparameters set with default values. The choice of this algorithm effectively translates into using L2 regularization.

**Figure 4.**
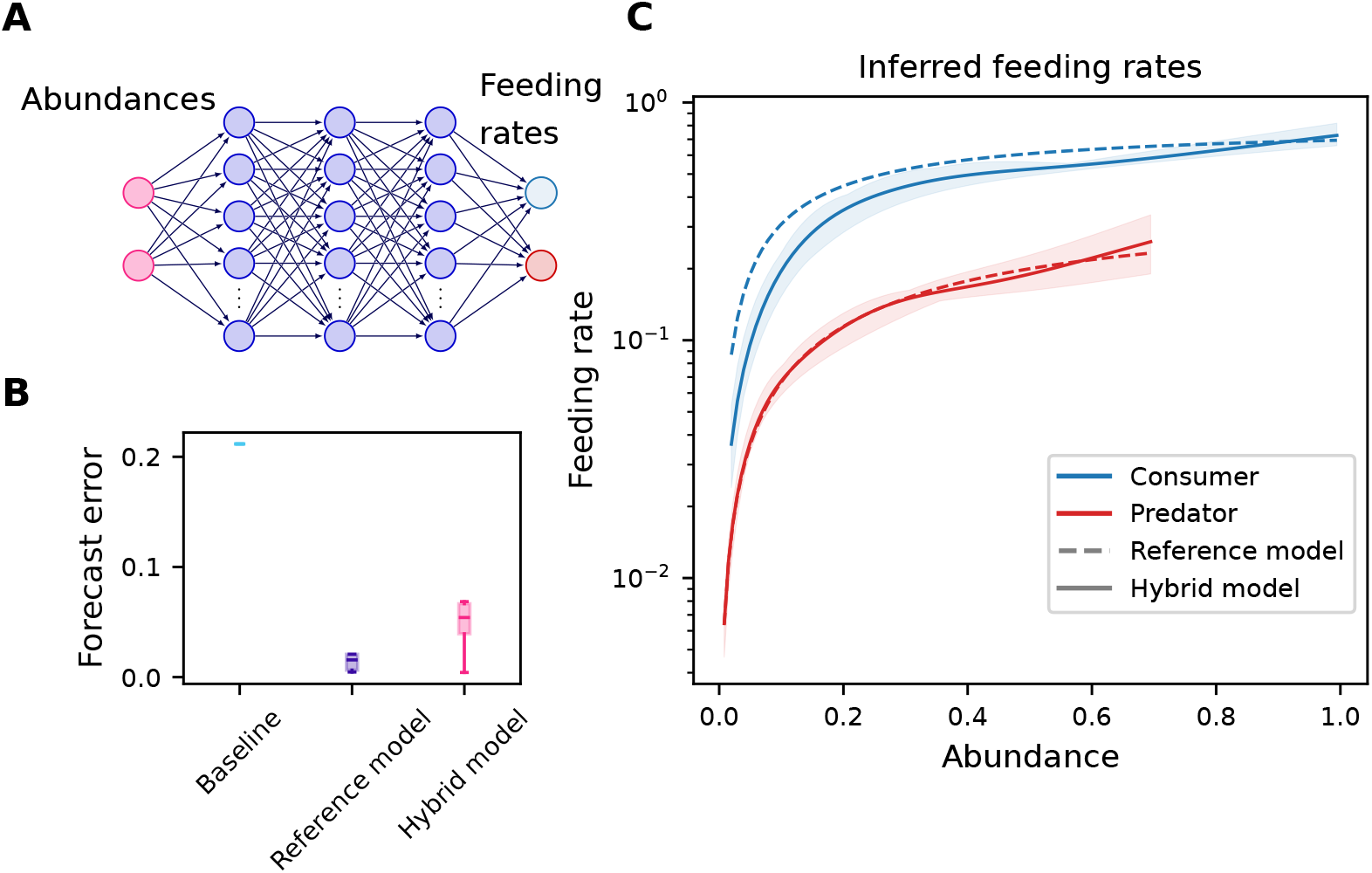
Neural network-based parameterization of feeding rates. **A**. Neural network architecture: the network takes as input the abundances of the primary producer and consumer species and predicts the per-capita feeding rates of the consumer and predator species. **B**. Forecast error of the trained hybrid model compared to that obtained from a baseline model with constant predictions (“Baseline”) and to the true generating model fitted under the same conditions for reference (“Reference model”). The hybrid model achieves similar forecast skill to the reference model, despite being more flexible. **C**. Post-hoc comparison of the neural network output against the true feeding rates (see Eq. (S1) and Eq. (S2) for details). The neural network successfully captures the functional form of the feeding rates. The solid line represents the median neural network output from 5 simulation runs, the shaded region indicates the minimum and maximum predicted values, and the dashed line corresponds to the ground truth. Results correspond to a noise level of *r* = 0.1.

Grid search results over various hyperparameters (Fig. S5) remain consistent with previous sections, and we find optimal performance with the following hyperparameters: *S* = 9, initial condition estimation, *γ* = 10^−2^, *λ* = 10^−9^, and *N*_*h*_ = 2^3^.

The trained hybrid model performs remarkably well, achieving forecast skill significantly better than a baseline using median training values for predictions (“Baseline” in Fig. 4**B**, *p <* 0.001 under Welch’s t-test) and similar to the true generating model ℳ_3_ fitted under identical conditions (“Reference model” in Fig. 4**B**, *p* = 0.141 under Welch’s test). Post-hoc comparison of inferred feeding rates with true functional responses (Fig. 4**C**) shows that the neural network successfully recovers their functional form, highlighting the effectiveness of hybrid modeling for feeding rates from time series.

We note that this scenario implicitly incorporates food web structure by setting per-capita feeding rates to zero except for consumer-producer and predator-consumer interactions. Additional inductive biases, such as separate neural networks for each functional response, could enhance interpretability and performance by explicitly encoding that feeding rates depend only on corresponding prey species.

#### 3.3.2 Neural network-based parameterization of the primary producer growth rate

We consider a variant of model ℳ_3_ where primary producer growth rate depends on a time-varying environmental forcing (e.g., temperature [Amarasekare and Johnson, 2017], see Eq. (S3)). We train a hybrid model where this growth rate is parameterized by a feed-forward neural network taking environmental forcing as input (Fig. 5**A**). The network architecture resembles that in Section 3.3.1 but with single input and output nodes. Grid search results, reported in Fig. S6, again show consistency with previous sections, yielding optimal performance with *S* = 9, initial condition estimation, *γ* = 10^−2^, *λ* = 10^−5^, and *N*_*h*_ = 2^2^.

**Figure 5.**
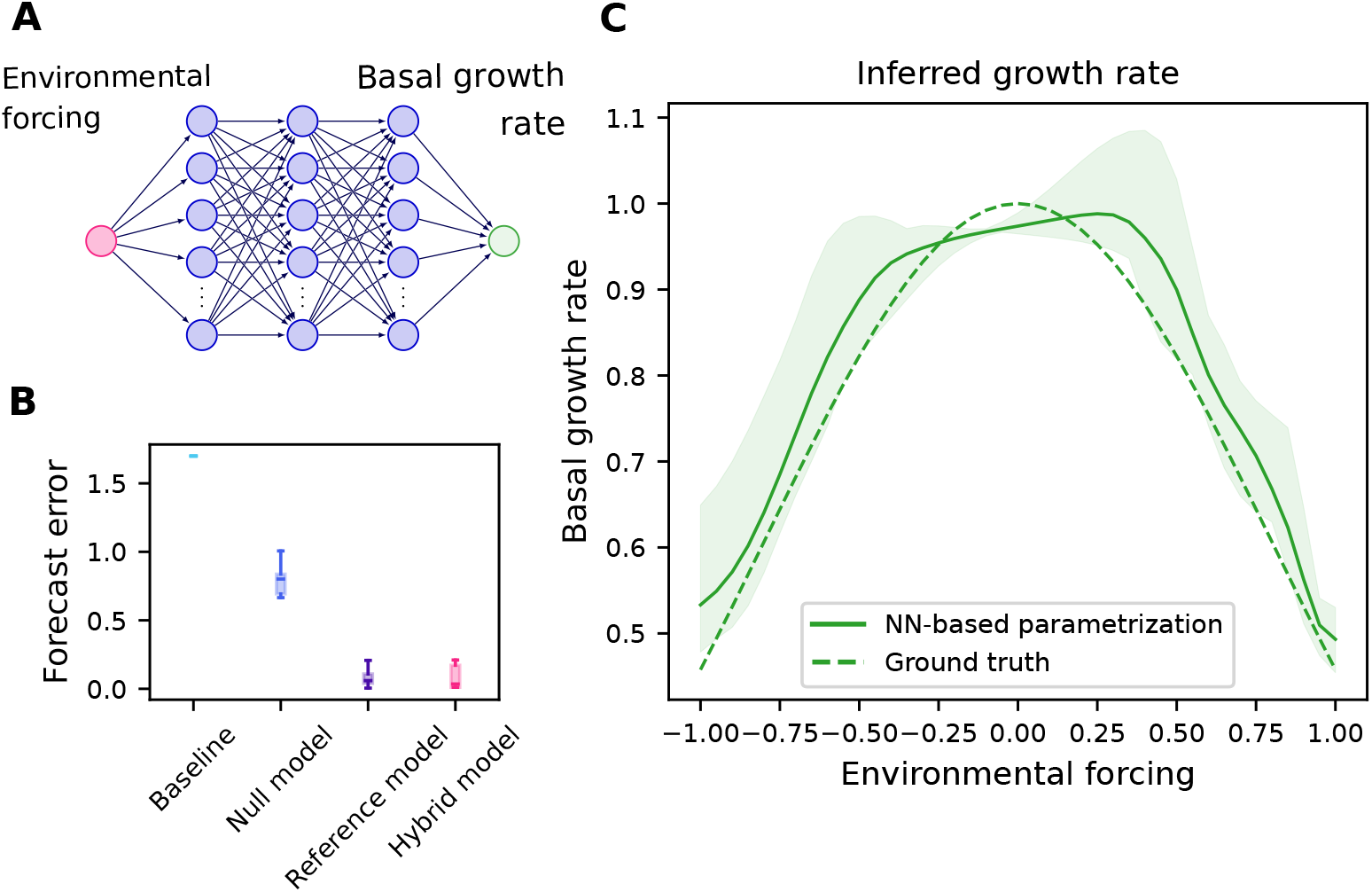
Neural network-based parameterization of the primary producer growth rate. **A**. Neural network architecture: the neural network takes the environmental forcing as input and predicts the basal growth rate of the primary producer species. **B**. Forecast error of the trained hybrid model compared to a baseline model with constant predictions (“Baseline”), a null model with a constant growth rate (“Null model”), and the true generating model fitted under the same conditions for reference (“Reference model”). The hybrid model performs significantly better than both the baseline and the null model (*p <* 0.001 under Welch’s t-test). **C**. Post-hoc comparison of the neural network output against the true primary producer growth rate from Eq. (S3) for 1*/s* = 1.2. The neural network accurately captures the dependence between the environmental forcing and the primary producer growth rate. The solid line represents the median neural network output from 10 simulation runs, the shaded regions indicate the minimum and maximum predicted values, and the dashed line corresponds to the ground truth. Results correspond to a noise level of *r* = 0.1.

The hybrid model’s forecast skill significantly exceeds both the baseline and a null model with constant growth rate (“Null model” in Fig. 5**B**, *p <* 0.001 under Welch’s t-test) and matches the performance of the true generating model fitted under identical conditions (*p* = 0.949 under Welch’s test, Fig. 5**B**). This demonstrates the framework’s capability for hypothesis testing, where comparing hybrid model performance against simpler models reveals the importance of time-varying environmental forcing. Post-hoc comparison between neural network output and true primary producer growth rate (Fig. 5**C**) shows accurate estimation of the underlying true Gaussian relationship, capturing both its mean and variance.

These results collectively demonstrate that our calibration framework successfully enables the construction of hybrid ecosystem models, where ecological processes with uncertain mathematical formulations can be effectively parameterized using neural networks while preserving mechanistic interpretability.

## 4 Discussion

We have developed a robust and scalable calibration framework for dynamic models that addresses fundamental challenges posed by ecological dynamics and ecological time series. Our piecewise formulation of the inverse problem, combined with independent estimation of initial conditions, substantially improves the convergence of gradient descent algorithms and Monte Carlo sampling methods when calibrating ecosystem models (Fig. 2). Furthermore, by automatically generating the gradient of the loss function required by gradient-based optimizers, differentiable programming and sensitivity analysis methods streamline the calibration process, enabling rapid model development cycles. Crucially, the robustness and scalability of our calibration framework enable hybrid modeling approaches, where neural networks parameterize complex ecological processes within mechanistic models. Such hybrid modeling could significantly improve forecasts of ecosystem dynamics while maintaining mechanistic interpretability, thereby helping to anticipate ecosystem responses to global changes [Urban et al., 2016].

Our work contributes to ongoing efforts to better assimilate observational data into mechanistic models [Schartau et al., 2017, Raissi et al., 2019, Kashinath et al., 2021, Auger-Méthé et al., 2021, Rosenbaum and Fronhofer, 2023, Paredes et al., 2023, Bolibar et al., 2023]. The segmentation strategy we propose builds upon multiple shooting methods [Pisarenko and Sornette, 2004, Hamilton, 2011], extending these approaches by integrating (i) a more flexible partitioning scheme with varying segment overlap, (ii) independent estimation of state variables, (iii) mini-batching capabilities, and (iv) integration with modern differentiable programming techniques [Sapienza et al., 2024]. While calibrating ordinary differential equation models by matching simulations against the full time series or using one-step-ahead forecasts are common practices in the ecological literature (see [Turchin, 2003, Rosenbaum and Fronhofer, 2023] and references therein), our benchmarks indicate that these strategies are suboptimal. Instead, selecting intermediate segment lengths significantly enhances both parameter estimation accuracy and forecast performance.

Collocation methods, such as those proposed by Yazdani et al. [2020], represent an interesting alternative approach to dynamic model calibration. These methods rely on parametric functions, typically neural networks, to emulate the full system dynamics. The parametric functions are trained to satisfy both observational data and additional constraints provided by the mechanistic model, which are incorporated through an additional loss function term weighted by a “goodness of fit” parameter. Although this approach can accommodate stochastic mechanistic models, it requires designing neural network architectures to approximate the complete system dynamics, thereby introducing additional hyperparameters that may negatively affect calibration and bias model selection [Ramsay et al., 2007]. In contrast, our framework uses neural networks to parameterize specific processes *within* the mechanistic model while directly matching model outputs against observational data. This approach leverages efficient differentiable programming methods to differentiate directly through forward model simulations, bypassing the need to design and fit neural networks that emulate full system dynamics. Consequently, our parameterization process becomes simpler, more interpretable, and more amenable to model selection. We note that while our work focused exclusively on differential equation-based models, we anticipate that the benefits of our calibration framework extend to discrete-time models also commonly used in ecology [Turchin, 2003]. HybridDynamicModels.jlreadily implements utilities for these types of models (see Section 2.2.2).

Our calibration framework addresses several practical constraints inherent to ecological datasets [Turchin, 2003]. By independently estimating state variables, the method accommodates incomplete and noisy observational data. Additionally, since segments are treated independently, the framework naturally extends to multiple independent time series. This capability is critical given that ecological surveys are typically shallow in time but composed of many spatial or temporal replicates [Hsieh et al., 2008, Clark et al., 2015, Ye and Sugihara, 2016, Pinsky et al., 2013, Dornelas et al., 2018, Burrows et al., 2019]. Moreover, the framework is particularly well-suited for model selection and hypothesis testing, enabling researchers to rapidly confront mechanistic models with empirical data [Curtsdotter et al., 2019, Becks et al., 2010]. The HybridDynamicModels.jllibrary further streamlines this process by providing composable components that can be easily recombined to test different model structures and parameterizations.

Key ecological processes in dynamic ecosystem models exhibit structural inaccuracies that have limited their predictive capabilities [Urban et al., 2016, Nadeau and Urban, 2019]. Hybrid modeling presents a promising solution that could improve the representation of these processes, enhancing the forecast skill of mechanistic models while preserving interpretability. Although hybrid approaches have been proposed in the context of ecological forecasting decades ago [Wood, 2001], their widespread adoption has been hindered by the daunting computational challenges of dynamic model calibration [Turchin, 2003]. Our library HybridDynamicModels.jlreduces the computational barriers that have previously limited its application. The transition from classical function approximations, such as the polynomials and splines proposed by Wood [2001], to neural networks opens up new possibilities for representing complex ecological processes. Neural networks excel in high-dimensional settings, where they can automatically construct higher-order features from multimodal data streams, making them particularly valuable for integrating a wealth of environmental covariates or structured data like satellite imagery. The possibility to incorporate multimodal environmental covariates into process-based models represents a significant opportunity for improving mechanistic forecasts of ecological dynamics.

Successful hybrid modeling requires careful consideration of several factors. Neural network design, including input selection, architecture, and output integration, must avoid collinearity with other model components to maintain the inductive bias of the model and its mechanistic interpretability. For instance, we found that restricting neural network parameterization to nonzero feeding rates is crucial when modeling functional responses. Before interpreting the outputs of the neural network, we recommend validating hybrid models against null models using cross-validation to confirm that learned representations are meaningful and supported by the data.

While our piecewise formulation of the inverse problem improves the convergence of inference methods, several limitations merit consideration. First, by reducing the curvature of the posterior distribution, the piecewise formulation introduces a loss of precision. We expect this limitation can be addressed through iterative training strategies that begin with short segments to identify probable parameter regions, then gradually increase segment length while reducing learning rates to refine estimates [Vilimelis Aceituno et al., 2025]. Second, parameter identifiability remains a fundamental challenge to maintain mechanistic interpretability, which is compromised when available data insufficiently constrain model behavior. Pre-experimental simulations using synthetic data can help assess sampling adequacy [Banks et al., 2017, Laubmeier et al., 2018], while model complexity may require reduction to align with the information content of available time series. Third, our current framework assumes constant parameter values across time series, yet real-world systems often exhibit temporal or spatial variability. This limitation can be addressed through partial pooling approaches [Beaumont, 2010] or by parameterizing model components in terms of covariates specific to different time series [Pahlow et al., 2008, Bolibar et al., 2023]. Finally, when parameter interpretation is crucial, combining our training strategy with Monte Carlo sampling methods provides comprehensive uncertainty quantification. However, these approaches scale poorly with the dimensionality of the parameter space, resulting in prohibitive computational costs for complex models. Approximate Bayesian inference methods, such as automatic differentiation variational inference [Kucukelbir et al., 2017], represent promising alternatives readily compatible with HybridDynamicModels.jlthrough its Turing.jlinterface [Fjelde et al., 2025], although we have not yet explored this avenue.

## 5 Conclusion

The rapid development of ecological monitoring technologies, including environmental DNA [Ruppert et al., 2019], remote sensing [Jetz et al., 2019], bioacoustics [Aide et al., 2013], and citizen science initiatives [GBIF: The Global Biodiversity Information Facility, 2022], is generating increasingly rich ecological time series. While this wealth of data has primarily benefited fully data-driven methods such as deep learning-based species distribution models [Gillespie et al., 2024], these approaches lack interpretability and robustness when extrapolating to novel environmental conditions. Our calibration framework has the potential to promote a more interpretable and robust approach to ecological forecasting through hybrid models. Looking ahead, the key challenge lies in designing models that embed appropriate inductive biases to balance predictive performance with interpretability while optimally leveraging available data. This requires careful consideration of which model components should be mechanistically hard-coded versus parameterized with neural networks. The development of standardized benchmarks could help guide the development of such hybrid approaches and accelerate progress [Picek et al., 2026]. We anticipate demonstrating applications to empirical ecosystem dynamics in the near future, with the ultimate goal of revealing the invariant mechanistic foundations underlying ecological dynamics.

## 6 Acknowledgements

L.P. and V.B. were supported by grants 205556 and 10005227 from the Swiss National Science Foundation. P.V.A. was supported by an ETH postdoctoral fellowship.

## 8 Code availability

The calibration framework is implemented in the Julia package HybridDynamicModels.jlavailable at https://github.com/vboussange/HybridDynamicModels.jl, and the simulation code for the experiments in this paper is available at https://github.com/vboussange/HybridDynamicModelExperiments.jl.

## 9 Contributions

**Conceptualization** V.B. and P.V.

**Formal Analysis** V.B., P.V. and F.S.

**Funding Acquisition** P.V. and L.P.

**Investigation** V.B. and P.V.

**Methodology** V.B. and F.S.

**Software** V.B. and F.S.

**Validation** V.B.

**Visualization** V.B.

**Supervision** L.P.

**Writing – Original Draft Preparation** V.B., P.V., F.S., and L.P.

## S1 Community model

### S1.1 Food-web ecosystem model

Denoting the abundance of species *i* by *N*_*i*_, and defining *N* = (*N*_1_, …, *N*_S_), the generalized community model can be expressed as

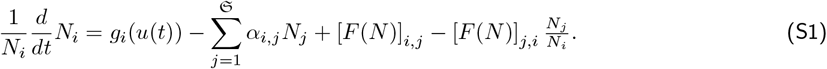

In Eq. (S1), the first two terms capture intrinsic growth rate and intra- and interspecific competition. The last two terms capture growth and loss due to trophic interactions. The per capita growth rate *g*_*i*_ may depend on a time-dependent environmental forcing *u*(*t*), but is set to a constant value unless specified. The competition coefficient between species *i* and *j* is denoted by *α*_*i,j*_. The per capita feeding rate [*F* (*N* )]_*i,j*_ of species *i* on *j* follows a functional response of type II

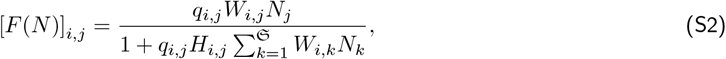

where *q*_*i,j*_ and *H*_*i,j*_ represent the attack rate and handling time of species *i* when feeding on species *j*, respectively, and *W*_*i,j*_ is the adjacency matrix of the feeding network (*W*_*i,j*_ = 1 if species *i* eats species *j* and 0 otherwise).

#### S1.1.1 Time-varying forcing three-species model variant

For the variant of *M*_3_ where the primary producer growth rate is depending on a forcing, we define *g*_1_ as:

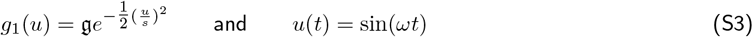

where 𝔤 is the maximum primary producer growth rate, reached when *u* = 0, and 1*/s* is the growth rate sensitivity to the forcing.

#### S1.1.2 Neural network-based parametrization of the per-capita feeding rates

For the hybrid model where the per-capita feeding rates are parametrized by a neural network, by denoting the neural network as NN: ℝ^2^ × *P* → ℝ^2^, we set [*F* (*N* )]_*i,j*_ in Eq. (S2) as

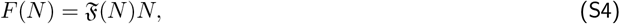

with ℱ (*N* ) being a matrix full of 0 except at [ℱ (*N* )]_2,1_ = [NN(*N*_1_, *N*_2_)]_1_ and at [ℱ (*N* )]_3,2_ = [NN(*N*_1_, *N*_2_)]_2_. This formulation permits to write the last term of Eq. (S2) as:

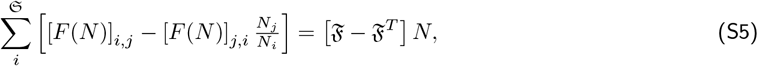

therefore avoiding a division by a state variable, which could lead to numerical problems.

### S1.2 Parameter value

ℳ_3_. We used the biologically realistic parameter values 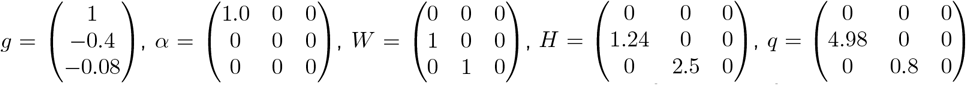, with initial conditions 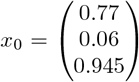. The dynamics of the system are chaotic for this set of parameter values [McCann and Yodzis, 1994a]. All non-zero coefficients were set as free parameters to be inferred, excluding the coefficients of the adjacency matrix *W* .

ℳ_5_. We used the biologically realistic parameter values 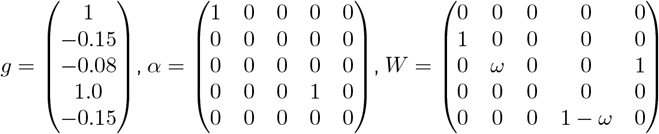 with 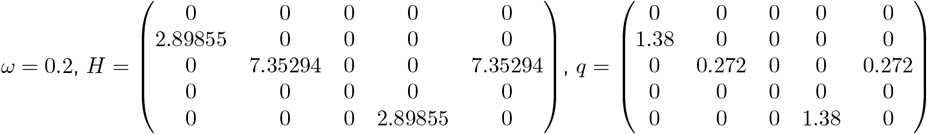, with initial conditions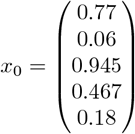 . The dynamics of the system are chaotic for this set of parameter values [Post et al., 2000]. All non-zero coefficients were set as free parameters, excluding the coefficients of the adjacency matrix *W*, where only *ω* was considered as a free parameter.

ℳ_7_. We used the biologically realistic parameter 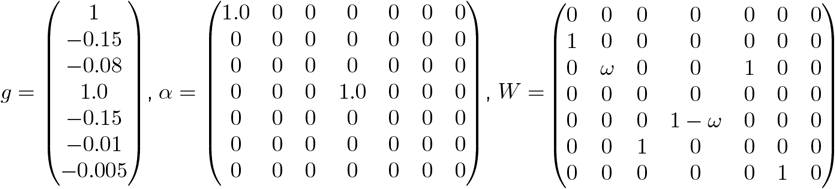 with values 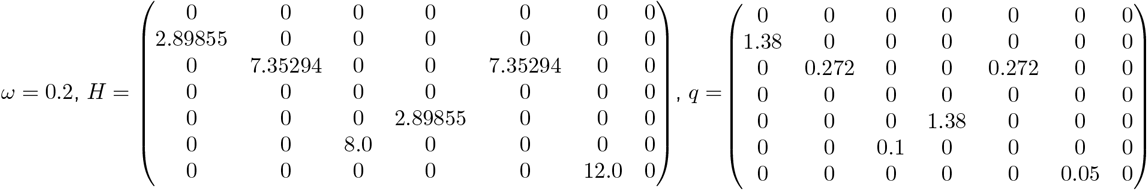, with initial conditions 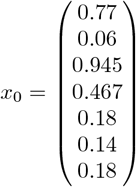. The dynamics of the system are chaotic for this set of parameter values. All non-zero coefficients were set as free parameters, excluding the coefficients of the adjacency matrix *W*, where only *ω* was considered as a free parameter.

#### Time-varying forcing three-species model variant

We used the same parameter values as for ℳ_3_ where g in the model variant is set to the same value as *g*_1_ in ℳ_3_, in addition to setting *ω* = 2*π/*3000.

## S2 Supplementary Figures

**Figure S1.**
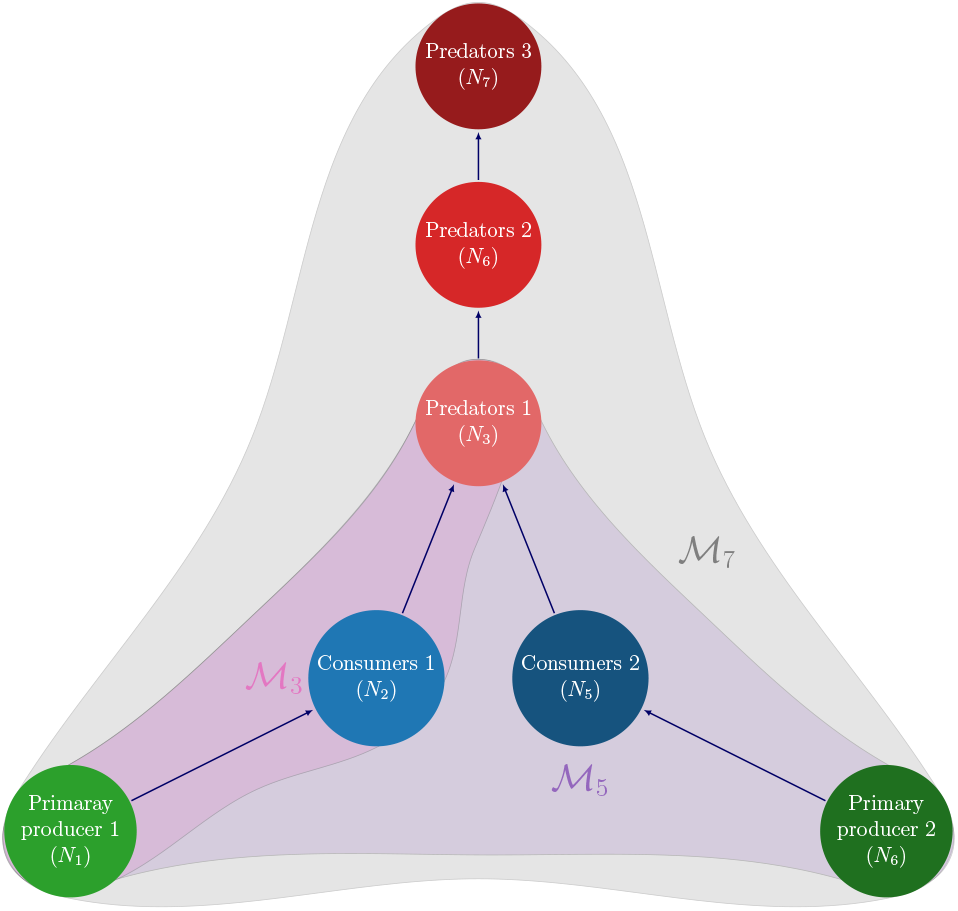
Food web systems considered. ℳ_3_ corresponds to the 3-species chaotic model presented in [Hastings and Powell, 1991], where a predator group feeds upon a consumer group, itself feeding upon on a primary producer group with type II functional responses. ℳ_5_ is an extension of ℳ_3_ proposed in [Post et al., 2000], where another food chain is linked to the foodweb through the predator group. ℳ_7_ extends ℳ_5_ by further considering two additional top predator groups.

**Figure S2.**
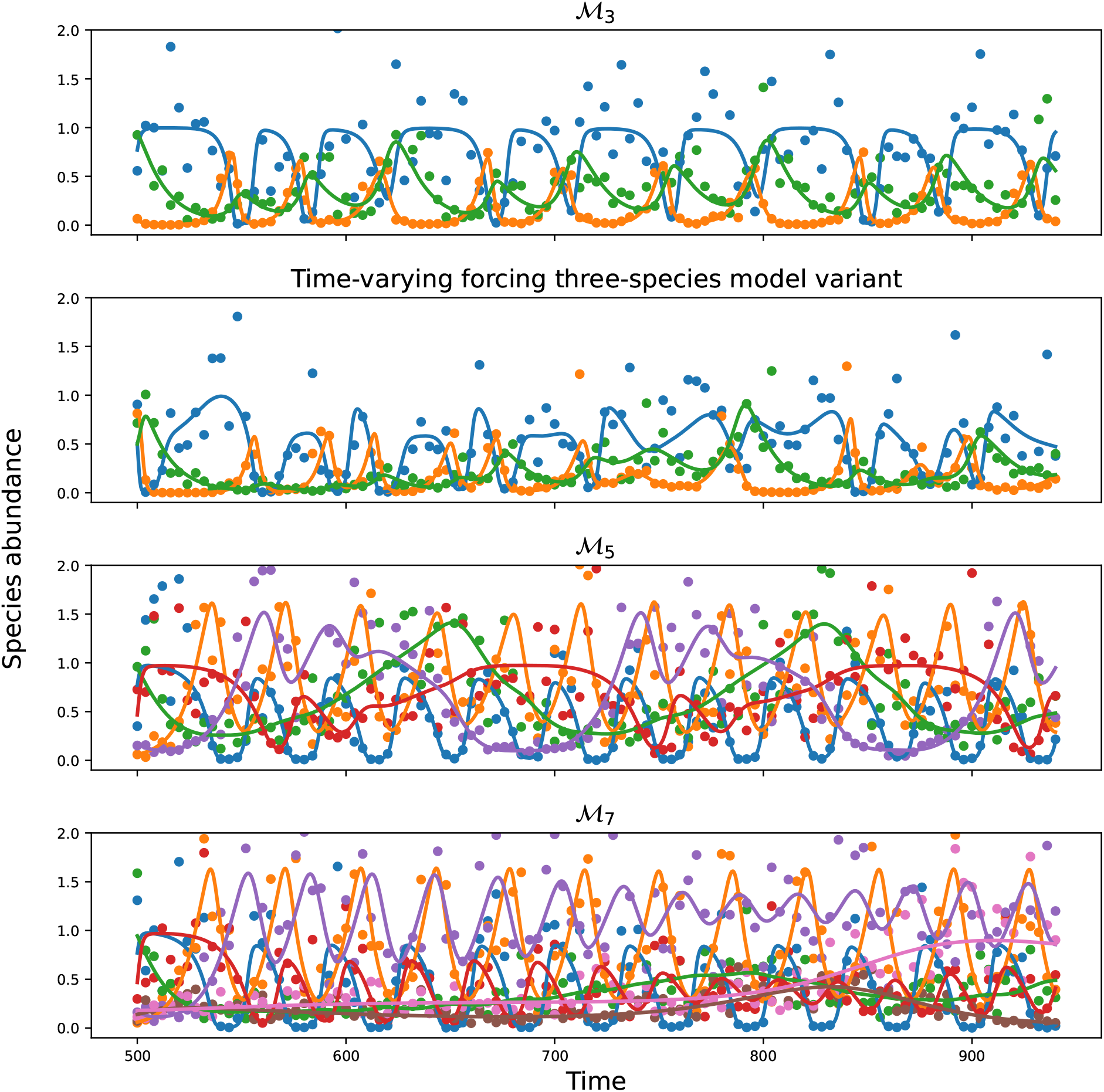
Training and test data obtained from the four community models and associated parameter values considered, generated with a noise level *r* = 0.4.

**Figure S3.**
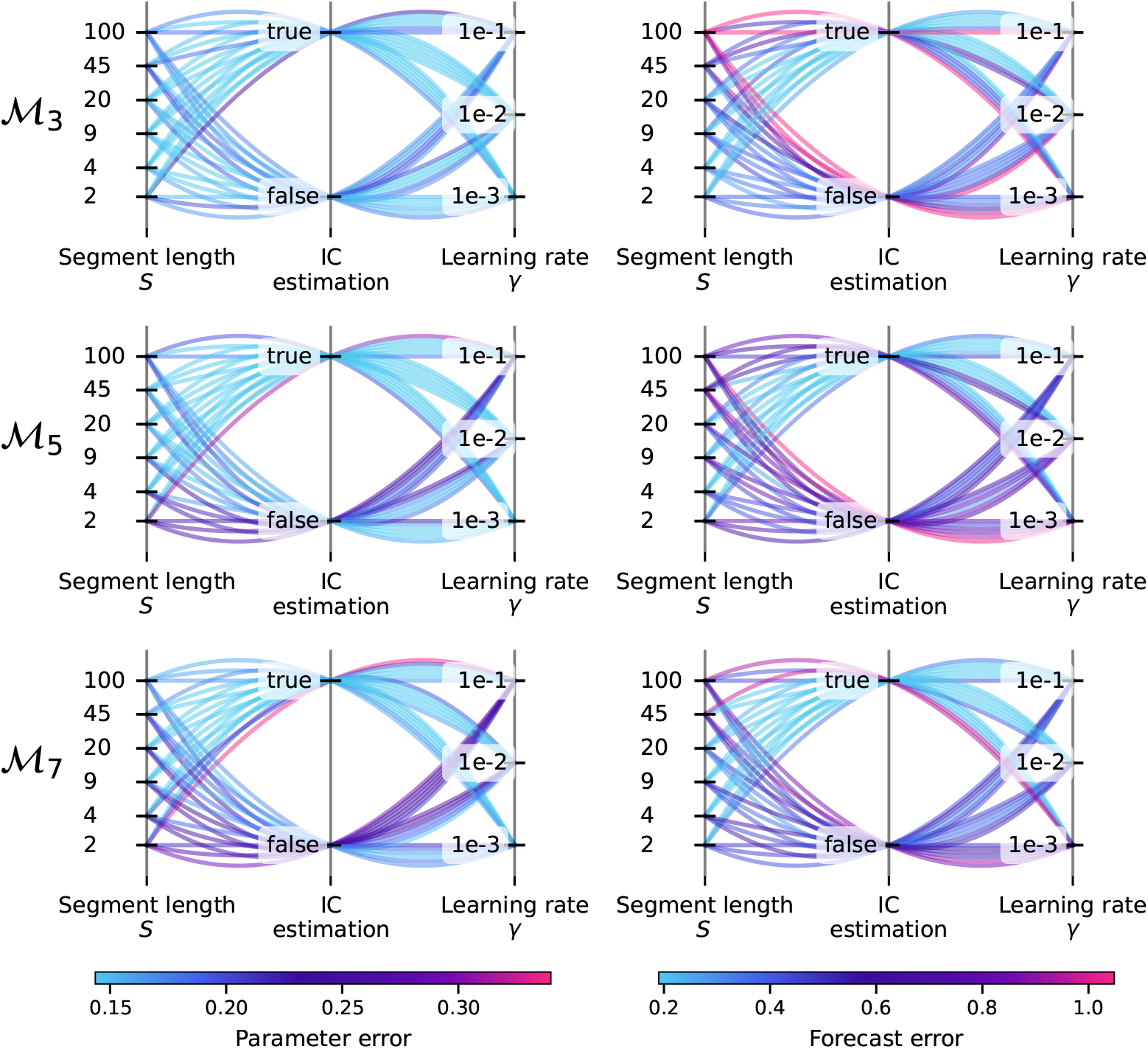
Parallel coordinates plot showing the effect of the combination of segment length *S*, estimation of ICs and learning rates on parameter error and forecast error, for varying food-web model complexity. Results are obtained with a perturbation parameter *ϵ* = 1, a noise level *r* = 0.4 a fixed minibatch size *b* = min(10, *M* ) and a segment shift *R* = *S* − 1.

**Figure S4.**
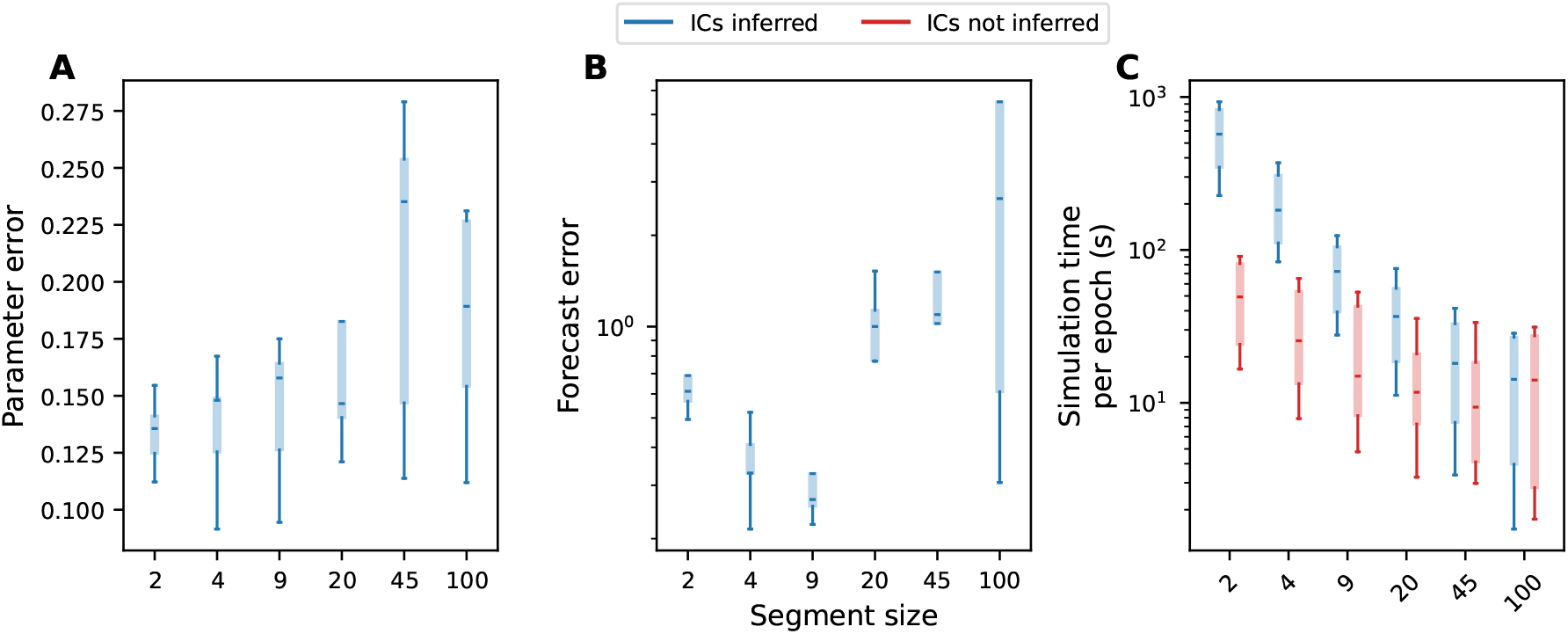
Effect of segment length *S* on parameter error, forecast error and simulation time, with model ℳ_3_ in a full posterior distribution estimation context with Hamiltonian Monte Carlo (HMC) sampling. We only report parameter and forecast error in the case where initial conditions are not inferred, as it proves to be too computationally expensive otherwise. **A**. Parameter error against segment length *S*. **B**. Forecast error against segment length *S*. **C**. Simulation time for a single iteration of the HMC algorithm against segment length *S*. In all panels, results are obtained from 5 simulation runs.

**Figure S5.**
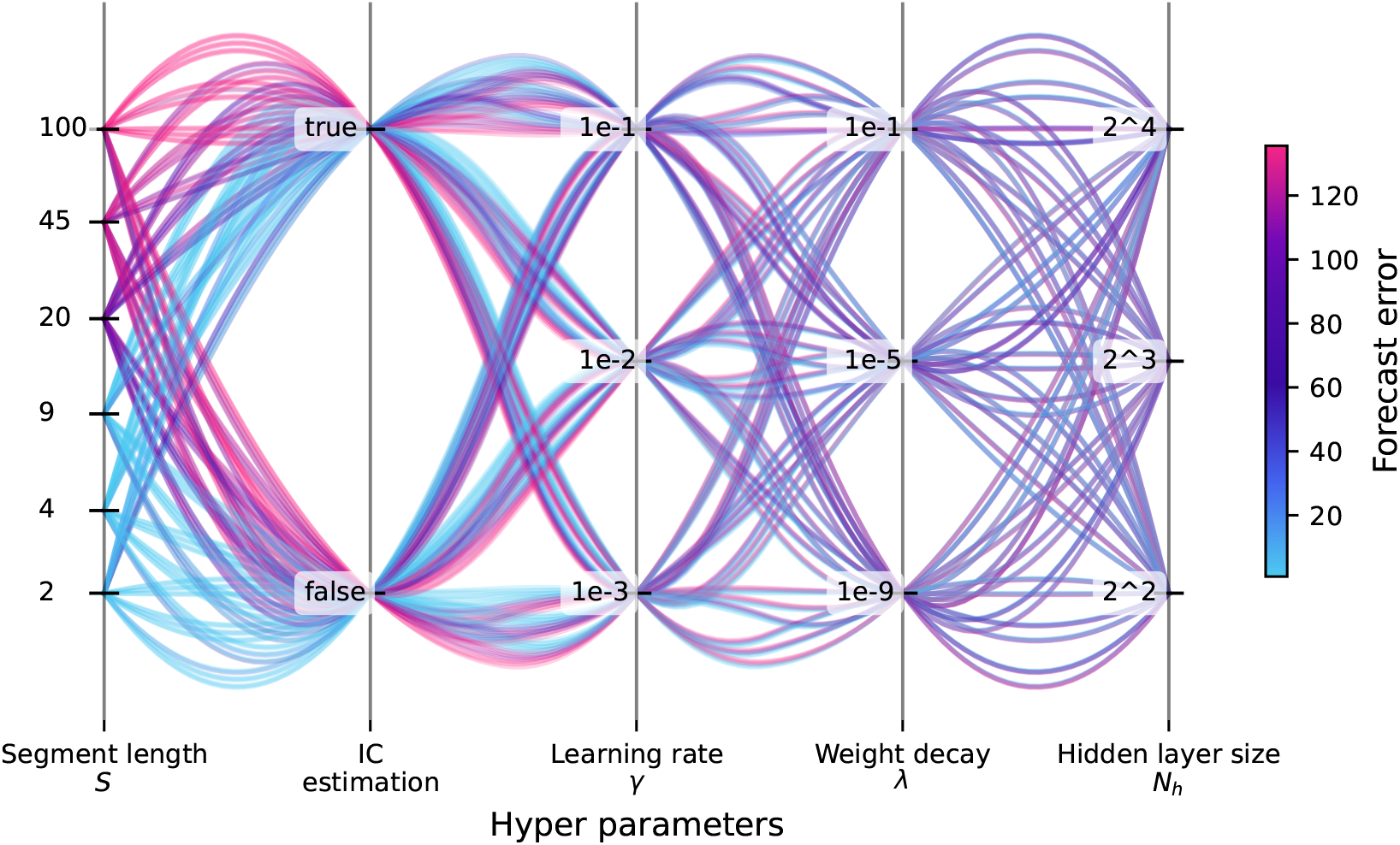
Parallel coordinates plot showing the effect of the various hyperparameters on the forecast error obtained with the hybrid model in Section 3.3.1. Results are obtained with a perturbation parameter *ϵ* = 1, a noise level *r* = 0.1 a fixed minibatch size *b* = min(10, *M* ) and a segment shift *R* = *S* + 1.

**Figure S6.**
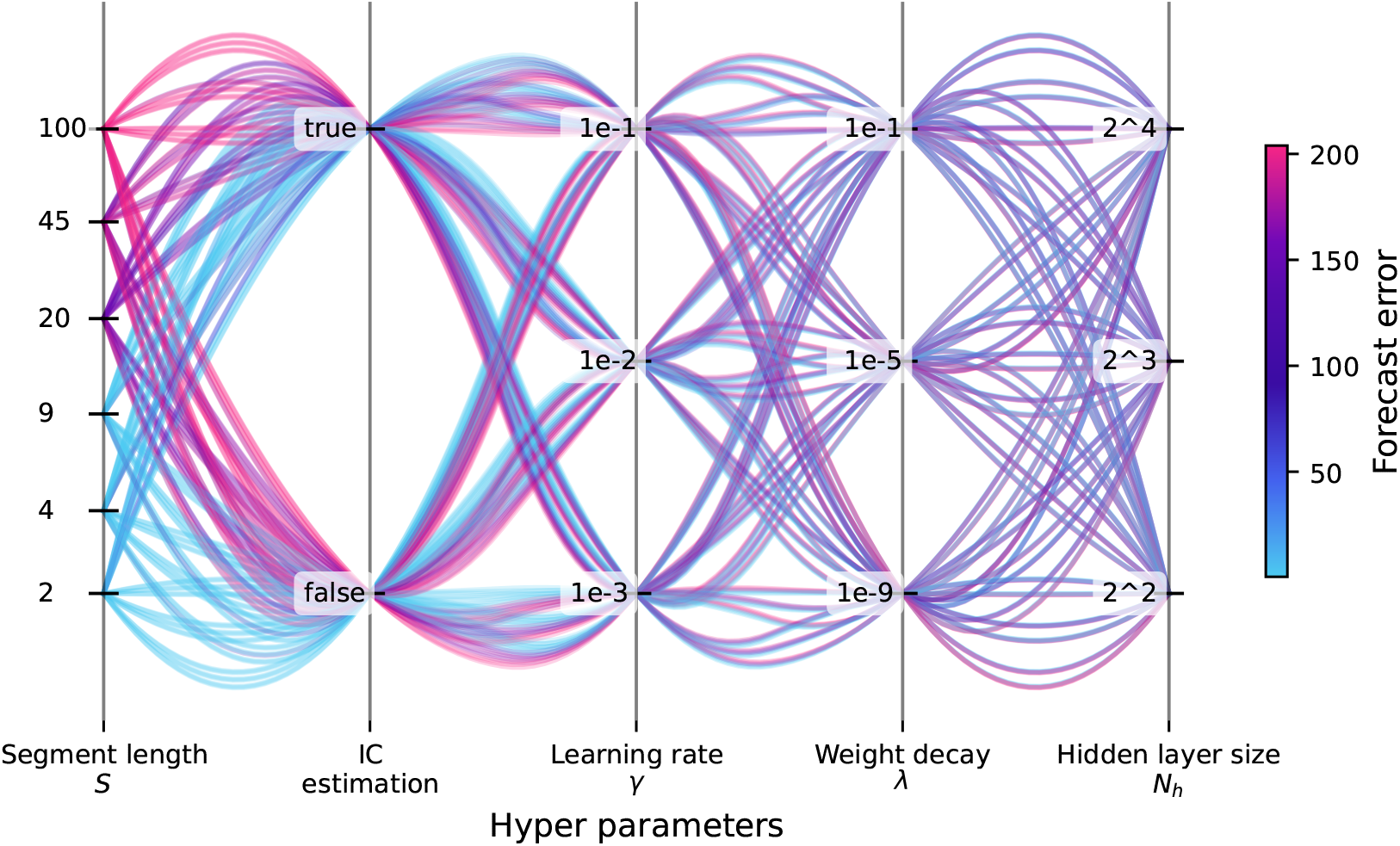
Parallel coordinates plot showing the effect of the various hyperparameters on the forecast error obtained with the hybrid model in Section 3.3.2. Results are obtained with a perturbation parameter *ϵ* = 1, a noise level *r* = 0.1 a fixed minibatch size *b* = min(10, *M* ) and a segment shift *R* = *S* + 1.

## References

T. Mitchell Aide, Carlos Corrada-Bravo, Marconi Campos-Cerqueira, Carlos Milan, Giovany Vega, and Rafael Alvarez. Real-time bioacoustics monitoring and automated species identification. PeerJ, 1(1):e103, July 2013. ISSN 2167-8359. doi: 10.7717/peerj.103.

Anna Åkesson, Alva Curtsdotter, Anna Eklöf, Bo Ebenman, Jon Norberg, and György Barabás. The importance of species interactions in eco-evolutionary community dynamics under climate change. Nature Communications, 12(1): 4759, December 2021. ISSN 2041-1723. doi: 10.1038/s41467-021-24977-x.

Mark Alber, Adrian Buganza Tepole, William R. Cannon, Suvranu De, Salvador Dura-Bernal, Krishna Garikipati, George Karniadakis, William W. Lytton, Paris Perdikaris, Linda Petzold, and Ellen Kuhl. Integrating machine learning and multiscale modeling—perspectives, challenges, and opportunities in the biological, biomedical, and behavioral sciences. npj Digital Medicine, 2(1):115, December 2019. ISSN 2398-6352. doi: 10.1038/s41746-019-0193-y.

Priyanga Amarasekare and Christopher Johnson. Evolution of Thermal Reaction Norms in Seasonally Varying Environments. The American Naturalist, 189(3):E31–E45, March 2017. ISSN 0003-0147, 1537-5323. doi: 10.1086/690293.

Marie Auger-Méthé, Ken Newman, Diana Cole, Fanny Empacher, Rowenna Gryba, Aaron A. King, Vianey Leos-Barajas, Joanna Mills Flemming, Anders Nielsen, Giovanni Petris, and Len Thomas. A guide to state–space modeling of ecological time series. Ecological Monographs, 91(4):e01470, 2021. ISSN 1557-7015. doi: 10.1002/ecm.1470.

H.T. Banks, J.E. Banks, R. Bommarco, A. Curtsdotter, T. Jonsson, and A.N. Laubmeier. PARAMETER ESTIMATION FOR AN ALLOMETRIC FOOD WEB MODEL. International Journal of Pure and Apllied Mathematics, 114 (1):143–160, April 2017. ISSN 1311-8080. doi: 10.12732/ijpam.v114i1.12.

Anthony D. Barnosky, Elizabeth A. Hadly, Jordi Bascompte, Eric L. Berlow, James H. Brown, Mikael Fortelius, Wayne M. Getz, John Harte, Alan Hastings, Pablo A. Marquet, Neo D. Martinez, Arne Mooers, Peter Roopnarine, Geerat Vermeij, John W. Williams, Rosemary Gillespie, Justin Kitzes, Charles Marshall, Nicholas Matzke, David P. Mindell, Eloy Revilla, and Adam B. Smith. Approaching a state shift in Earth’s biosphere. Nature, 486(7401): 52–58, 2012. ISSN 00280836. doi: 10.1038/nature11018.

Mark A. Beaumont. Approximate Bayesian Computation in Evolution and Ecology. Annual Review of Ecology, Evolution, and Systematics, 41(1): 379–406, 2010. ISSN 1543-592X. doi: 10.1146/annurev-ecolsys-102209-144621.

Lutz Becks, Stephen P. Ellner, Laura E. Jones, and Jr G. Hairston Nelson G. Reduction of adaptive genetic diversity radically alters eco-evolutionary community dynamics. Ecology Letters, 13(8):989–997, May 2010. ISSN 14610248. doi: 10.1111/j.1461-0248.2010.01490.x.

Elisa Benincà, Jef Huisman, Reinhard Heerkloss, Klaus D. Jöhnk, Pedro Branco, Egbert H. Van Nes, Marten Scheffer, and Stephen P. Ellner. Chaos in a long-term experiment with a plankton community. Nature, 451(7180):822–825, February 2008. ISSN 0028-0836. doi: 10.1038/nature06512.

Ottar N. Bjørnstad and Bryan T. Grenfell. Noisy Clockwork: Time Series Analysis of Population Fluctuations in Animals. Science, 293(5530):638–643, July 2001. ISSN 0036-8075. doi: 10.1126/science.1062226.

Jordi Bolibar, Facundo Sapienza, Fabien Maussion, Redouane Lguensat, Bert Wouters, and Fernando Pérez. Universal differential equations for glacier ice flow modelling. Geoscientific Model Development, 16(22):6671–6687, November 2023. ISSN 1991-959X. doi: 10.5194/gmd-16-6671-2023.

Ian L. Boyd. The Art of Ecological Modeling. Science, 337(6092):306–307, July 2012. ISSN 0036-8075. doi: 10.1126/science.1225049.

Natalie J. Briscoe, Jane Elith, Roberto Salguero-Gómez, José J. Lahoz-Monfort, James S. Camac, Katherine M. Giljohann, Matthew H. Holden, Bronwyn A. Hradsky, Michael R. Kearney, Sean M. McMahon, Ben L. Phillips, Tracey J. Regan, Jonathan R. Rhodes, Peter A. Vesk, Brendan A. Wintle, Jian D.L. Yen, and Gurutzeta Guillera-Arroita. Forecasting species range dynamics with process-explicit models: Matching methods to applications. Ecology Letters, 22(11): 1940–1956, 2019. ISSN 1461-0248. doi: 10.1111/ele.13348.

Michael T. Burrows, Amanda E. Bates, Mark J. Costello, Martin Edwards, Graham J. Edgar, Clive J. Fox, Benjamin S. Halpern, Jan G. Hiddink, Malin L. Pinsky, Ryan D. Batt, Jorge García Molinos, Benjamin L. Payne, David S. Schoeman, Rick D. Stuart-Smith, and Elvira S. Poloczanska. Ocean community warming responses explained by thermal affinities and temperature gradients. Nature Climate Change, 9(12):959–963, December 2019. ISSN 1758-678X. doi: 10.1038/s41558-019-0631-5.

Juliano Sarmento Cabral, Luis Valente, and Florian Hartig. Mechanistic simulation models in macroecology and biogeography: State-of-art and prospects. Ecography, 40(2): 267–280, 2017. ISSN 1600-0587. doi: 10.1111/ecog.02480.

Loïc Chalmandrier, Daniel B. Stouffer, Adam S. T. Purcell, William G. Lee, Andrew J. Tanentzap, and Daniel C. Laughlin. Predictions of biodiversity are improved by integrating trait-based competition with abiotic filtering. Ecology Letters, 25(5): 1277–1289, 2022. ISSN 1461-0248. doi: 10.1111/ele.13980.

Adam Thomas Clark, Hao Ye, Forest Isbell, Ethan R. Deyle, Jane Cowles, G. David Tilman, George Sugihara, and B. D. Inouye. Spatial convergent cross mapping to detect causal relationships from short time series. Ecology, 96 (5):1174–1181, 2015. ISSN 00129658. doi: 10.1890/14-1479.1.

Alva Curtsdotter, H. Thomas Banks, John E. Banks, Mattias Jonsson, Tomas Jonsson, Amanda N. Laubmeier, Michael Traugott, and Riccardo Bommarco. Ecosystem function in predator–prey food webs—confronting dynamic models with empirical data. Journal of Animal Ecology, 88(2):196–210, February 2019. ISSN 0021-8790. doi: 10.1111/1365-2656.12892.

Donald L. DeAngelis and Simeon Yurek. Equation-free modeling unravels the behavior of complex ecological systems. Proceedings of the National Academy of Sciences, 112(13):3856–3857, March 2015. ISSN 0027-8424. doi: 10.1073/pnas.1503154112.

John P. DeLong, Torrance C. Hanley, and David A. Vasseur. Predator-prey dynamics and the plasticity of predator body size. Functional Ecology, 28(2):487–493, April 2014. ISSN 02698463. doi: 10.1111/1365-2435.12199.

Benjamin Deneu, Maximilien Servajean, Pierre Bonnet, Christophe Botella, François Munoz, and Alexis Joly. Convolutional neural networks improve species distribution modelling by capturing the spatial structure of the environment. PLOS Computational Biology, 17(4):e1008856, April 2021. ISSN 1553-7358. doi: 10.1371/journal.pcbi.1008856.

Ethan R. Deyle, Robert M. May, Stephan B. Munch, and George Sugihara. Tracking and forecasting ecosystem interactions in real time. Proceedings of the Royal Society B: Biological Sciences, 283(1822):20152258, January 2016. ISSN 0962-8452. doi: 10.1098/rspb.2015.2258.

Maria Dornelas, Laura H. Antão, Faye Moyes, Amanda E. Bates, Anne E. Magurran, Dušan Adam, Asem A. Akhmetzhanova, Ward Appeltans, José Manuel Arcos, Haley Arnold, Narayanan Ayyappan, Gal Badihi, Andrew H. Baird, Miguel Barbosa, Tiago Egydio Barreto, Claus Bässler, Alecia Bellgrove, Jonathan Belmaker, Lisandro Benedetti-Cecchi, Brian J. Bett, Anne D. Bjorkman, Magdalena Błażewicz, Shane A. Blowes, Christopher P. Bloch, Timothy C. Bonebrake, Susan Boyd, Matt Bradford, Andrew J. Brooks, James H. Brown, Helge Bruelheide, Phaedra Budy, Fernando Carvalho, Edward Castañeda-Moya, Chaolun Allen Chen, John F. Chamblee, Tory J. Chase, Laura Siegwart Collier, Sharon K. Collinge, Richard Condit, Elisabeth J. Cooper, J. Hans C. Cornelissen, Unai Cotano, Shannan Kyle Crow, Gabriella Damasceno, Claire H. Davies, Robert A. Davis, Frank P. Day, Steven Degraer, Tim S. Doherty, Timothy E. Dunn, Giselda Durigan, J. Emmett Duffy, Dor Edelist, Graham J. Edgar, Robin Elahi, Sarah C. Elmendorf, Anders Enemar, S. K. Morgan Ernest, Rubén Escribano, Marc Estiarte, Brian S. Evans, Tung-Yung Fan, Fabiano Turini Farah, Luiz Loureiro Fernandes, Fábio Z. Farneda, Alessandra Fidelis, Robert Fitt, Anna Maria Fosaa, Geraldo Antonio Daher Correa Franco, Grace E. Frank, William R. Fraser, Hernando García, Roberto Cazzolla Gatti, Or Givan, Elizabeth Gorgone-Barbosa, William A. Gould, Corinna Gries, Gary D. Grossman, Julio R. Gutierréz, Stephen Hale, Mark E. Harmon, John Harte, Gary Haskins, Donald L. Henshaw, Luise Hermanutz, Pamela Hidalgo, Pedro Higuchi, Andrew Hoey, Gert Van Hoey, Annika Hofgaard, Kristen Holeck, Robert D. Hollister, Richard Holmes, Mia Hoogenboom, Chih-hao Hsieh, Stephen P. Hubbell, Falk Huettmann, Christine L. Huffard, Allen H. Hurlbert, Natália Macedo Ivanauskas, David Janík, Ute Jandt, Anna Jażdżewska, Tore Johannessen, Jill Johnstone, Julia Jones, Faith A. M. Jones, Jungwon Kang, Tasrif Kartawijaya, Erin C. Keeley, Douglas A. Kelt, Rebecca Kinnear, Kari Klanderud, Halvor Knutsen, Christopher C. Koenig, Alessandra R. Kortz, Kamil Král, Linda A. Kuhnz, Chao-Yang Kuo, David J. Kushner, Claire Laguionie-Marchais, Lesley T. Lancaster, Cheol Min Lee, Jonathan S. Lefcheck, Esther Lévesque, David Lightfoot, Francisco Lloret, John D. Lloyd, Adrià López-Baucells, Maite Louzao, Joshua S. Madin, BorgÞór Magnússon, Shahar Malamud, Iain Matthews, Kent P. McFarland, Brian McGill, Diane McKnight, William O. McLarney, Jason Meador, Peter L. Meserve, Daniel J. Metcalfe, Christoph F. J. Meyer, Anders Michelsen, Nataliya Milchakova, Tom Moens, Even Moland, Jon Moore, Carolina Mathias Moreira, Jörg Müller, Grace Murphy, Isla H. Myers-Smith, Randall W. Myster, Andrew Naumov, Francis Neat, James A. Nelson, Michael Paul Nelson, Stephen F. Newton, Natalia Norden, Jeffrey C. Oliver, Esben M. Olsen, Vladimir G. Onipchenko, Krzysztof Pabis, Robert J. Pabst, Alain Paquette, Sinta Pardede, David M. Paterson, Raphaël Pélissier, Josep Peñuelas, Alejandro Pérez-Matus, Oscar Pizarro, Francesco Pomati, Eric Post, Herbert H. T. Prins, John C. Priscu, Pieter Provoost, Kathleen L. Prudic, Erkki Pulliainen, B. R. Ramesh, Olivia Mendivil Ramos, Andrew Rassweiler, Jose Eduardo Rebelo, Daniel C. Reed, Peter B. Reich, Suzanne M. Remillard, Anthony J. Richardson, J. Paul Richardson, Itai van Rijn, Ricardo Rocha, Victor H. Rivera-Monroy, Christian Rixen, Kevin P. Robinson, Ricardo Ribeiro Rodrigues, Denise de Cerqueira Rossa-Feres, Lars Rudstam, Henry Ruhl, Catalina S. Ruz, Erica M. Sampaio, Nancy Rybicki, Andrew Rypel, Sofia Sal, Beatriz Salgado, Flavio A. M. Santos, Ana Paula Savassi-Coutinho, Sara Scanga, Jochen Schmidt, Robert Schooley, Fakhrizal Setiawan, Kwang-Tsao Shao, Gaius R. Shaver, Sally Sherman, Thomas W. Sherry, Jacek Siciński, Caya Sievers, Ana Carolina da Silva, Fernando Rodrigues da Silva, Fabio L. Silveira, Jasper Slingsby, Tracey Smart, Sara J. Snell, Nadejda A. Soudzilovskaia, Gabriel B. G. Souza, Flaviana Maluf Souza, Vinícius Castro Souza, Christopher D. Stallings, Rowan Stanforth, Emily H. Stanley, José Mauro Sterza, Maarten Stevens, Rick Stuart-Smith, Yzel Rondon Suarez, Sarah Supp, Jorge Yoshio Tamashiro, Sukmaraharja Tarigan, Gary P. Thiede, Simon Thorn, Anne Tolvanen, Maria Teresa Zugliani Toniato, Ørjan Totland, Robert R. Twilley, Gediminas Vaitkus, Nelson Valdivia, Martha Isabel Vallejo, Thomas J. Valone, Carl Van Colen, Jan Vanaverbeke, Fabio Venturoli, Hans M. Verheye, Marcelo Vianna, Rui P. Vieira, Tomáš Vrška, Con Quang Vu, Lien Van Vu, Robert B. Waide, Conor Waldock, Dave Watts, Sara Webb, Tomasz Wesołowski, Ethan P. White, Claire E. Widdicombe, Dustin Wilgers, Richard Williams, Stefan B. Williams, Mark Williamson, Michael R. Willig, Trevor J. Willis, Sonja Wipf, Kerry D. Woods, Eric J. Woehler, Kyle Zawada, Michael L. Zettler, and Thomas Hickler. BioTIME: A database of biodiversity time series for the Anthropocene. Global Ecology and Biogeography, 27(7):760–786, July 2018. ISSN 1466-822X. doi: 10.1111/geb.12729.

Katja Fennel, Martin Losch, Jens Schröter, and Manfred Wenzel. Testing a marine ecosystem model: Sensitivity analysis and parameter optimization. Journal of Marine Systems, 28(1–2):45–63, February 2001. ISSN 09247963. doi: 10.1016/S0924-7963(00)00083-X.

J. Fiechter, R. Herbei, W. Leeds, J. Brown, R. Milliff, C. Wikle, A. Moore, and T. Powell. A Bayesian parameter estimation method applied to a marine ecosystem model for the coastal Gulf of Alaska. Ecological Modelling, 258: 122–133, June 2013. ISSN 03043800. doi: 10.1016/j.ecolmodel.2013.03.003.

Rosie A. Fisher, Charles D. Koven, William R.L. Anderegg, Bradley O. Christoffersen, Michael C. Dietze, Caroline E. Farrior, Jennifer A. Holm, George C. Hurtt, Ryan G. Knox, Peter J. Lawrence, Jeremy W. Lichstein, Marcos Longo, Ashley M. Matheny, David Medvigy, Helene C. Muller-Landau, Thomas L. Powell, Shawn P. Serbin, Hisashi Sato, Jacquelyn K. Shuman, Benjamin Smith, Anna T. Trugman, Toni Viskari, Hans Verbeeck, Ensheng Weng, Chonggang Xu, Xiangtao Xu, Tao Zhang, and Paul R. Moorcroft. Vegetation demographics in Earth System Models: A review of progress and priorities. Global Change Biology, 24(1): 35–54, 2018. ISSN 13652486. doi: 10.1111/gcb.13910.

Tor Erlend Fjelde, Kai Xu, David Widmann, Mohamed Tarek, Cameron Pfiffer, Martin Trapp, Seth D. Axen, Xianda Sun, Markus Hauru, Penelope Yong, Will Tebbutt, Zoubin Ghahramani, and Hong Ge. Turing.jl: A general-purpose probabilistic programming language. 2025. doi: 10.1145/3711897. URL https://doi.org/10.1145/3711897.

Damien A. Fordham, Cleo Bertelsmeier, Barry W. Brook, Regan Early, Dora Neto, Stuart C. Brown, Sébastien Ollier, and Miguel B. Araújo. How complex should models be? Comparing correlative and mechanistic range dynamics models. Global Change Biology, 24(3): 1357–1370, 2018. ISSN 1365-2486. doi: 10.1111/gcb.13935.

GBIF: The Global Biodiversity Information Facility. What is GBIF? 2022.

William L. Geary, Michael Bode, Tim S. Doherty, Elizabeth A. Fulton, Dale G. Nimmo, Ayesha I. T. Tulloch, Vivitskaia J. D. Tulloch, and Euan G. Ritchie. A guide to ecosystem models and their environmental applications. Nature Ecology & Evolution, 4(11):1459–1471, November 2020. ISSN 2397-334X. doi: 10.1038/s41559-020-01298-8.

M. Gehlen, R. Barciela, L. Bertino, P. Brasseur, M. Butenschön, F. Chai, A. Crise, Y. Drillet, D. Ford, D. Lavoie,P. Lehodey, C. Perruche, A. Samuelsen, and E. Simon. Building the capacity for forecasting marine biogeochemistry and ecosystems: Recent advances and future developments. Journal of Operational Oceanography, 8(sup1):s168–s187, April 2015. ISSN 1755-876X. doi: 10.1080/1755876X.2015.1022350.

Wendy Gentleman, Andrew Leising, Bruce Frost, Suzanne Strom, and James Murray. Functional responses for zooplankton feeding on multiple resources: A review of assumptions and biological dynamics. Deep Sea Research Part II: Topical Studies in Oceanography, 50(22–26):2847–2875, November 2003. ISSN 09670645. doi: 10.1016/j.dsr2.2003.07.001.

Lauren E. Gillespie, Megan Ruffley, and Moises Exposito-Alonso. Deep learning models map rapid plant species changes from citizen science and remote sensing data. 121(37):e2318296121, 2024. doi: 10.1073/pnas.2318296121. URL https://www.pnas.org/doi/full/10.1073/pnas.2318296121.

Varun Godbole, George E. Dahl, Justin Gilmer, Christopher J. Shallue, and Zachary Nado. Deep learning tuning playbook, 2023. URL http://github.com/google-research/tuning_playbook.

Sanmitra Gosh, Paul Birrell, and Daniela De Angelis. Variational inference for nonlinear ordinary differential equations. Proceedings of The 24th International Conference on Artificial Intelligence and Statistics, 130(29): 2719–2727, 2021.

Antoine Guisan and Niklaus E. Zimmermann. Predictive habitat distribution models in ecology. Ecological Modelling, 135(2):147–186, December 2000. ISSN 0304-3800. doi: 10.1016/S0304-3800(00)00354-9.

Ryan N. Gutenkunst, Joshua J. Waterfall, Fergal P. Casey, Kevin S. Brown, Christopher R. Myers, and James P. Sethna. Universally Sloppy Parameter Sensitivities in Systems Biology Models. PLoS Computational Biology, 3(10): e189, October 2007. ISSN 1553-7358. doi: 10.1371/journal.pcbi.0030189.

Franz Hamilton. Parameter Estimation in Differential Equations: A Numerical Study of Shooting Methods. 4: 16–31, 2011. ISSN 23277807. doi: 10.1137/10S010739. URL http://www.siam.org/students/siuro/vol4/S01073.pdf.

Florian Hartig, James Dyke, Thomas Hickler, Steven I. Higgins, Robert B. O’Hara, Simon Scheiter, and Andreas Huth. Connecting dynamic vegetation models to data - an inverse perspective: Dynamic vegetation models – an inverse perspective. Journal of Biogeography, 39(12):2240–2252, December 2012. ISSN 03050270. doi: 10.1111/j.1365-2699.2012.02745.x.

Alan Hastings and Thomas Powell. Chaos in a Three-Species Food Chain. Ecology, 72(3):896–903, June 1991. ISSN 00129658. doi: 10.2307/1940591.

Alan Hastings, Carole L. Hom, Stephen Ellner, Peter Turchin, and H. Charles J. Godfray. Chaos in Ecology: Is Mother Nature a Strange Attractor? Annual Review of Ecology and Systematics, 24(1):1–33, November 1993. ISSN 0066-4162. doi: 10.1146/annurev.es.24.110193.000245.

Lukas Heiland, Georges Kunstler, Vladimír Šebeň, and Lisa Hülsmann. Which demographic processes control competitive equilibria? Bayesian calibration of a size-structured forest population model. Ecology and Evolution, 13(7): e10232, 2023. ISSN 2045-7758. doi: 10.1002/ece3.10232.

Steven I. Higgins, Simon Scheiter, and Mahesh Sankaran. The stability of African savannas: Insights from the indirect estimation of the parameters of a dynamic model. Ecology, 91(6):1682–1692, June 2010. ISSN 0012-9658. doi: 10.1890/08-1368.1.

Matthew D. Hoffman and Andrew Gelman. The No-U-Turn Sampler: Adaptively Setting Path Lengths in Hamiltonian Monte Carlo. 2011. doi: 10.48550/arXiv.1111.4246. URL http://arxiv.org/abs/1111.4246.

Chih-hao Hsieh, Christian Anderson, and George Sugihara. Extending Nonlinear Analysis to Short Ecological Time Series. The American Naturalist, 171(1):71–80, January 2008. ISSN 0003-0147. doi: 10.1086/524202.

Jef Huisman and Franz J. Weissing. Biodiversity of plankton by species oscillations and chaos. Nature, 402(6760): 407–410, November 1999. ISSN 0028-0836. doi: 10.1038/46540.

IPBES. Summary for policymakers of the global assessment report on biodiversity and ecosystem services. Technical report, Zenodo, November 2019.

Walter Jetz, Melodie A. McGeoch, Robert Guralnick, Simon Ferrier, Jan Beck, Mark J. Costello, Miguel Fernandez, Gary N. Geller, Petr Keil, Cory Merow, Carsten Meyer, Frank E. Muller-Karger, Henrique M. Pereira, Eugenie C. Regan, Dirk S. Schmeller, and Eren Turak. Essential biodiversity variables for mapping and monitoring species populations. Nature Ecology and Evolution, 3(4): 539–551, 2019. ISSN 2397334X. doi: 10.1038/s41559-019-0826-1.

K. Kashinath, M. Mustafa, A. Albert, J. L. Wu, C. Jiang, S. Esmaeilzadeh, K. Azizzadenesheli, R. Wang, A. Chattopadhyay, A. Singh, A. Manepalli, D. Chirila, R. Yu, R. Walters, B. White, H. Xiao, H. A. Tchelepi, P. Marcus, A. Anandkumar, P. Hassanzadeh, and Prabhat. Physics-informed machine learning: Case studies for weather and climate modelling. Philosophical Transactions of the Royal Society A: Mathematical, Physical and Engineering Sciences, 379(2194), 2021. ISSN 1364503X. doi: 10.1098/rsta.2020.0093.

Diederik P. Kingma and Jimmy Ba. Adam: A Method for Stochastic Optimization. pages 1–15, December 2014.

Aaron Klebanoff and Alan Hastings. Chaos in three species food chains. Journal of Mathematical Biology, 32(5): 427–451, May 1994. ISSN 0303-6812. doi: 10.1007/BF00160167.

Alp Kucukelbir, Dustin Tran, Rajesh Ranganath, Andrew Gelman, and David M. Blei. Automatic differentiation variational inference. Journal of Machine Learning Research, 18(14): 1–45, 2017. URL http://jmlr.org/papers/v18/16-107.html.

A. N. Laubmeier, Kate Wootton, J. E. Banks, Riccardo Bommarco, Alva Curtsdotter, Tomas Jonsson, Tomas Roslin, and H. T. Banks. From theory to experimental design—Quantifying a trait-based theory of predator-prey dynamics. PLOS ONE, 13(4):e0195919, April 2018. ISSN 1932-6203. doi: 10.1371/journal.pone.0195919.

Risto Lignell, Heikki Haario, Marko Laine, and T. Frede Thingstad. Getting the “right” parameter values for models of the pelagic microbial food web. Limnology and Oceanography, 58(1):301–313, January 2013. ISSN 00243590. doi: 10.4319/lo.2013.58.1.0301.

Ilya Loshchilov and Frank Hutter. Decoupled Weight Decay Regularization, 2019. URL http://arxiv.org/abs/1711.05101.

Yingbo Ma, Vaibhav Dixit, Michael J. Innes, Xingjian Guo, and Chris Rackauckas. A Comparison of Automatic Differentiation and Continuous Sensitivity Analysis for Derivatives of Differential Equation Solutions. In 2021 IEEE High Performance Extreme Computing Conference (HPEC), number 2, pages 1–9. IEEE, September 2021. ISBN 978-1-66542-369-4. doi: 10.1109/HPEC49654.2021.9622796.

Richard J. Matear. Parameter optimization and analysis of ecosystem models using simulated annealing: A case study at Station P. Journal of Marine Research, 53(4):571–607, July 1995. ISSN 00222402. doi: 10.1357/0022240953213098.

Kevin McCann and Alan Hastings. Re–evaluating the omnivory–stability relationship in food webs. Proceedings of the Royal Society of London. Series B: Biological Sciences, 264(1385):1249–1254, August 1997. ISSN 0962-8452. doi: 10.1098/rspb.1997.0172.

Kevin McCann and Peter Yodzis. Biological Conditions for Chaos in a Three-Species Food Chain. Ecology, 75(2): 561–564, March 1994a. ISSN 00129658. doi: 10.2307/1939558.

Kevin McCann and Peter Yodzis. Nonlinear Dynamics and Population Disappearances. The American Naturalist, 144(5):873–879, November 1994b. ISSN 0003-0147. doi: 10.1086/285714.

Jonas M Mikhaeil, Zahra Monfared, and Daniel Durstewitz. On the difficulty of learning chaotic dynamics with RNNs. 2022.

Christopher P. Nadeau and Mark C. Urban. Eco-evolution on the edge during climate change. Ecography, 42(7): 1280–1297, 2019. ISSN 1600-0587. doi: 10.1111/ecog.04404.

Jon Norberg, Mark C. Urban, Mark Vellend, Christopher A. Klausmeier, and Nicolas Loeuille. Eco-evolutionary responses of biodiversity to climate change. Nature Climate Change, 2(10):747–751, October 2012. ISSN 1758-678X. doi: 10.1038/nclimate1588.

Jörn Pagel and Frank M. Schurr. Forecasting species ranges by statistical estimation of ecological niches and spatial population dynamics. Global Ecology and Biogeography, 21(2): 293–304, 2012. ISSN 1466822X. doi: 10.1111/j.1466-8238.2011.00663.x.

Markus Pahlow, Alain F. Vézina, Benoit Casault, Heidi Maass, Louise Malloch, Daniel G. Wright, and Youyu Lu. Adaptive model of plankton dynamics for the North Atlantic. Progress in Oceanography, 76(2): 151–191, 2008. ISSN 00796611. doi: 10.1016/j.pocean.2007.11.001.

Avik Pal. Lux: Explicit parameterization of deep neural networks in julia. 2023. doi: 10.5281/zenodo.7808903. URL https://doi.org/10.5281/zenodo.7808903.

Josefa Arán Paredes, Koen Hufkens, and Benjamin D. Stocker. Rsofun: A model-data integration framework for simulating ecosystem processes, November 2023.

Joanna S. Pelc, Ehouarn Simon, Laurent Bertino, Ghada El Serafy, and Arnold W. Heemink. Application of model reduced 4D-Var to a 1D ecosystem model. Ocean Modelling, 57–58:43–58, November 2012. ISSN 14635003. doi: 10.1016/j.ocemod.2012.09.003.

Grace C. Y. Peng, Mark Alber, Adrian Buganza Tepole, William R. Cannon, Suvranu De, Savador Dura-Bernal, Krishna Garikipati, George Karniadakis, William W. Lytton, Paris Perdikaris, Linda Petzold, and Ellen Kuhl. Multiscale Modeling Meets Machine Learning: What Can We Learn? Archives of Computational Methods in Engineering, 28 (3):1017–1037, May 2021. ISSN 1134-3060. doi: 10.1007/s11831-020-09405-5.

Charles T. Perretti, George Sugihara, and Stephan B. Munch. Nonparametric forecasting outperforms parametric methods for a simulated multispecies system. Ecology, 94(4):794–800, April 2013. ISSN 0012-9658. doi: 10.1890/12-0904.1.

Lukáš Picek, Stefan Kahl, Hervé Goëau, Lukáš Adam, Théo Larcher, Cesar Leblanc, Maximilien Servajean, Klára Janoušková, Jiří Matas, Vojtěch Čermák, Kostas Papafitsoros, Robert Planqué, Willem-Pier Vellinga, Holger Klinck, Tom Denton, Juan Sebastián Cañas, Giulio Martellucci, Fabrice Vinatier, Pierre Bonnet, and Alexis Joly. Overview of LifeCLEF 2025: Challenges on Species Presence Prediction and Identification, and Individual Animal Identification. In Jorge Carrillo-de-Albornoz, Alba García Seco de Herrera, Julio Gonzalo, Laura Plaza, Josiane Mothe, Florina Piroi, Paolo Rosso, Damiano Spina, Guglielmo Faggioli, and Nicola Ferro, editors, Experimental IR Meets Multilinguality, Multimodality, and Interaction, pages 338–362, Cham, 2026. Springer Nature Switzerland. ISBN 978-3-032-04354-2.

M. L. Pinsky, B. Worm, M. J. Fogarty, J. L. Sarmiento, and S. A. Levin. Marine Taxa Track Local Climate Velocities. Science, 341(6151):1239–1242, September 2013. ISSN 0036-8075. doi: 10.1126/science.1239352.

V. F. Pisarenko and D. Sornette. Statistical methods of parameter estimation for deterministically chaotic time series. Physical Review E, 69(3):036122, March 2004. ISSN 1539-3755. doi: 10.1103/PhysRevE.69.036122.

David M. Post, M. Elizabeth Conners, and Debra S. Goldberg. Prey preference by a top predator and the stability of linked food chains. Ecology, 81(1): 8–14, 2000. ISSN 00129658. doi: 10.1890/0012-9658(2000)081[0008:PPBATP]2.0.CO;2.

Drew Purves, Jörn P. W. Scharlemann, Mike Harfoot, Tim Newbold, Derek P. Tittensor, Jon Hutton, and Stephen Emmott. Time to model all life on Earth. Nature, 493(7432):295–297, January 2013. ISSN 0028-0836. doi: 10.1038/493295a.

Christopher Rackauckas and Qing Nie. DifferentialEquations.jl – A Performant and Feature-Rich Ecosystem for Solving Differential Equations in Julia. Journal of Open Research Software, 5, 2017. ISSN 2049-9647. doi: 10.5334/jors.151.

M. Raissi, P. Perdikaris, and G.E. Karniadakis. Physics-informed neural networks: A deep learning framework for solving forward and inverse problems involving nonlinear partial differential equations. Journal of Computational Physics, 378:686–707, February 2019. ISSN 00219991. doi: 10.1016/j.jcp.2018.10.045.

J. O. Ramsay, G. Hooker, D. Campbell, and J. Cao. Parameter estimation for differential equations: A generalized smoothing approach. Journal of the Royal Statistical Society: Series B (Statistical Methodology), 69(5):741–796, November 2007. ISSN 13697412. doi: 10.1111/j.1467-9868.2007.00610.x.

Stephan Rasp, Michael S. Pritchard, and Pierre Gentine. Deep learning to represent subgrid processes in climate models. Proceedings of the National Academy of Sciences, 115(39):9684–9689, September 2018. doi: 10.1073/pnas.1810286115.

Benjamin Rosenbaum and Emanuel A. Fronhofer. Confronting population models with experimental microcosm data: From trajectory matching to state-space models. Ecosphere, 14(4):e4503, April 2023. ISSN 2150-8925, 2150-8925. doi: 10.1002/ecs2.4503.

Benjamin Rosenbaum and Björn C. Rall. Fitting functional responses: Direct parameter estimation by simulating differential equations. Methods in Ecology and Evolution, 9(10):2076–2090, October 2018. ISSN 2041-210X. doi: 10.1111/2041-210X.13039.

Benjamin Rosenbaum, Michael Raatz, Guntram Weithoff, Gregor F. Fussmann, and Ursula Gaedke. Estimating Parameters From Multiple Time Series of Population Dynamics Using Bayesian Inference. Frontiers in Ecology and Evolution, 6(JAN), January 2019. ISSN 2296-701X. doi: 10.3389/fevo.2018.00234.

Krista M. Ruppert, Richard J. Kline, and Md Saydur Rahman. Past, present, and future perspectives of environmental DNA (eDNA) metabarcoding: A systematic review in methods, monitoring, and applications of global eDNA. Global Ecology and Conservation, 17:e00547, January 2019. ISSN 23519894. doi: 10.1016/j.gecco.2019.e00547.

Facundo Sapienza, Jordi Bolibar, Frank Schäfer, Brian Groenke, Avik Pal, Victor Boussange, Patrick Heimbach, Giles Hooker, Fernando Pérez, Per-Olof Persson, and Christopher Rackauckas. Differentiable Programming for Differential Equations: A Review. 2024. doi: 10.48550/arXiv.2406.09699. URL http://arxiv.org/abs/2406.09699.

Markus Schartau, Philip Wallhead, John Hemmings, Ulrike Löptien, Iris Kriest, Shubham Krishna, Ben A. Ward, Thomas Slawig, and Andreas Oschlies. Reviews and syntheses: Parameter identification in marine planktonic ecosystem modelling. Biogeosciences, 14(6):1647–1701, March 2017. ISSN 1726-4189. doi: 10.5194/bg-14-1647-2017.

Marten Scheffer, Steve Carpenter, Jonathan A Foley, Carl Folke, and Brian Walker. Catastrophic shifts in ecosystems. Nature, 413(6856):591–596, October 2001. ISSN 0028-0836. doi: 10.1038/35098000.

Simon Scheiter, Liam Langan, and Steven I. Higgins. Next-generation dynamic global vegetation models: Learning from community ecology. New Phytologist, 198(3):957–969, May 2013. ISSN 0028-646X. doi: 10.1111/nph.12210.

Y.H. Spitz, J.R. Moisan, M.R. Abbott, and J.G. Richman. Data assimilation and a pelagic ecosystem model: Parameterization using time series observations. Journal of Marine Systems, 16(1–2):51–68, September 1998. ISSN 09247963. doi: 10.1016/S0924-7963(97)00099-7.

Peter Turchin. Complex Population Dynamics. Princeton University Press, 2003. ISBN 978-0-691-09021-4. URL http://www.jstor.org/stable/j.ctt24hqkz.

M. C. Urban, G. Bocedi, A. P. Hendry, J.-B. Mihoub, G. Pe’er, A. Singer, J. R. Bridle, L. G. Crozier, L. De Meester, W. Godsoe, A. Gonzalez, J. J. Hellmann, R. D. Holt, A. Huth, K. Johst, C. B. Krug, P. W. Leadley, S. C. F. Palmer, J. H. Pantel, A. Schmitz, P. A. Zollner, and J. M. J. Travis. Improving the forecast for biodiversity under climate change. Science, 353(6304), September 2016. ISSN 0036-8075. doi: 10.1126/science.aad8466.

Pau Vilimelis Aceituno, Jack William Miller, Noah Marti, Youssef Farag, and Victor Boussange. Temporal horizons in forecasting: a performance-learnability trade-off. Manuscript submitted for publication, 2025.

Ben A. Ward, Marjorie A.M. Friedrichs, Thomas R. Anderson, and Andreas Oschlies. Parameter optimisation techniques and the problem of underdetermination in marine biogeochemical models. Journal of Marine Systems, 81 (1-2):34–43, April 2010. ISSN 09247963. doi: 10.1016/j.jmarsys.2009.12.005.

Jared Willard, Xiaowei Jia, Shaoming Xu, Michael Steinbach, and Vipin Kumar. Integrating Scientific Knowledge with Machine Learning for Engineering and Environmental Systems. 1(1): 1–35, 2020. ISSN 2331-8422.

Simon N. Wood. PARTIALLY SPECIFIED ECOLOGICAL MODELS. 71(1): 1–25, 2001. ISSN 0012-9615. doi: 10.1890/0012-9615(2001)071[0001:PSEM]2.0.CO;2. URL http://doi.wiley.com/10.1890/0012-9615(2001)071[0001:PSEM]2.0.CO;2.

Yongjin Xiao and Marjorie A. M. Friedrichs. The assimilation of satellite-derived data into a one-dimensional lower trophic level marine ecosystem model. Journal of Geophysical Research: Oceans, 119(4):2691–2712, April 2014. ISSN 2169-9275. doi: 10.1002/2013JC009433.

Tao Xu, Luther White, Dafeng Hui, and Yiqi Luo. Probabilistic inversion of a terrestrial ecosystem model: Analysis of uncertainty in parameter estimation and model prediction. Global Biogeochemical Cycles, 20(2):n/a–n/a, June 2006. ISSN 08866236. doi: 10.1029/2005GB002468.

Alireza Yazdani, Lu Lu, Maziar Raissi, and George Em Karniadakis. Systems biology informed deep learning for inferring parameters and hidden dynamics. PLOS Computational Biology, 16(11):e1007575, November 2020. ISSN 1553-7358. doi: 10.1371/journal.pcbi.1007575.

Hao Ye and George Sugihara. Information leverage in interconnected ecosystems: Overcoming the curse of dimensionality. Science, 353(6302):922–925, August 2016. ISSN 0036-8075. doi: 10.1126/science.aag0863.

Hao Ye, Richard J. Beamish, Sarah M. Glaser, Sue C. H. Grant, Chih-hao Hsieh, Laura J. Richards, Jon T. Schnute, and George Sugihara. Equation-free mechanistic ecosystem forecasting using empirical dynamic modeling. Proceedings of the National Academy of Sciences, 112(13):E1569–E1576, March 2015. ISSN 0027-8424. doi: 10.1073/pnas.1417063112.

Qing Zhu and Qianlai Zhuang. Ecosystem biogeochemistry model parameterization: Do more flux data result in a better model in predicting carbon flux? Ecosphere, 6(12):art283, December 2015. ISSN 2150-8925. doi: 10.1890/ES15-00259.1.

